# Elevated viral recombination in short-lived hosts

**DOI:** 10.1101/2025.10.01.679867

**Authors:** Mete Yuksel, Qiqi Yang, Matt Osmond, Nicole Mideo

**Author notes:** Equal contribution (Matt Osmond, Nicole Mideo).

## Abstract

Recombination (including reassortment) is a salient force in viral evolution and has been implicated in the emergence of several zoonotic pathogens in humans. Viral recombination occurs during simultaneous infection of an individual host with multiple genotypes (co-infection). Thus, processes which affect the incidence of a disease in the host population affect how often viral genotypes recombine. We investigate whether and how host traits affect the realized rate of viral recombination using a mathematical model that makes feedbacks between viral evolution and host ecology (in particular, lifespan) explicit. Our main result is that viruses of host species that are short-lived tend to recombine more frequently than those of relatively long-lived hosts. This is because of differences in population density and, thus, the prevalence of (co-)infection at equilibrium. Using highly pathogenic avian influenza sequence data, we test the prediction that recombination is elevated in short-lived hosts. In agreement with this prediction, the magnitude of statistical associations between mutations on different segments of the flu genome increases with host body size, a proxy for lifespan. Similarly, estimates of the reassortment rate from phylogenetic network analyses decrease with body size. We discuss the implications of these findings for disease emergence.

**Subject category:** Evolution

**Subject areas:** evolution, health and disease and epidemiology, ecology

## 1 Introduction

Recombination, i.e., the exchange of genetic information between different individuals or lineages, occurs in all domains of life and is central to understanding the evolution of sexual reproduction (1). A unique feature of recombination in viruses is that it only occurs when multiple virions co-infect the same host cell. Upon co-infection, recombination can take place when the viral polymerase dissociates from one template and completes transcription on another (2; 3; 4). Similarly, in segmented viruses, a process called reassortment can give rise to offspring with mixed ancestry when physically unlinked segments from co-infecting genotypes are packaged together (5; 6; 7). In either case, the constraint that virions must co-infect the same cell gives rise to stark differences in the rate of recombination between viral populations and species (2; 3; 8; 9; 10). These differences are the result of the mechanistic coupling of co-infection and recombination: processes that affect the former affect the latter.

Coupling of co-infection and viral recombination is consequential because recombination (inclusive of reassortment) has been implicated in the process by which pathogens infect and sustain transmission in new hosts (i.e., emergence; (4; 11; 12; 13; 14)) and is therefore important for human and animal health. In particular, there is phylogenetic evidence that intra- and inter-subtype reassortment occur frequently in the evolutionary history of influenza, and that re-assortments — involving gene segments normally circulating in birds, pigs, and humans — have driven all human influenza A pandemics after 1918 (6; 15). The SARS-like coronaviruses from which SARS-CoV-2 descends also have a reticulate evolutionary history; present-day sequence data suggest that recombination is elevated around the Spike open reading frame, and that many recombination events have occurred in the recent past (14; 16). Although there is disagreement about the importance of recombination events in the spillover and subsequent spread of SARS-CoV-2 (17), this is compelling evidence that recombination fuels emergence in certain contexts.

The relationship between viral recombination and co-infection suggests that host ecology should influence how often viral genomes recombine and, in turn, how viral evolution affects disease emergence. Indeed, previous studies (18; 19; 20; 21) have found that certain host traits (e.g., body size) are strong predictors of zoonotic potential (e.g., the proportion of viruses in a given species capable of infecting humans). Using these associations, some have argued that bats are “special” reservoirs due to their longevity, elevated internal body temperatures, lack of apparent signs of disease, species richness, etc. (19; 22; 23; 24). Others have argued that rodents may be special because of their fast-paced life histories, marked by frequent reproduction, early age of sexual maturity, and rapid population turnover (21; 25). Other work has found that there are no differences in zoonotic risk between host taxa after controlling for reservoir species richness and viral richness (26). Using mathematical models, Nusimer et al. (27) recently brought some clarity to this debate. They found that (1) in the absence of viral evolution, short-lived hosts (e.g., rodents) are more likely to fuel spillover and emergence than ones that are relatively long-lived (e.g., bats) and (2) if there is a genetic trade-off between virus transmission in the reservoir host and in humans, the frequency of alleles favouring emergence is greatest when hosts are long-lived (27). These findings suggest that additional genetic details, such as the rate and consequences of viral recombination, matter for comparing the potential of reservoir taxa to harbor zoonotic diseases.

Here, we build on Nusimer et al. (27) to ask: in what reservoir host taxa (distinguished by mean lifespan, duration of infection, and the rate of waning immunity) is the rate of viral recombination highest? By analyzing a mathematical model that integrates feedbacks between viral recombination and co-infection, we generate predictions for how recombination rate scales with host traits. Our key result is that, due to differences in population density and co-infection, viral recombination is elevated in short-lived hosts compared to hosts that are relatively long-lived. We test this prediction using highly pathogenic avian influenza (HPAI) sequence data, sampled from different bird species, in two ways. First, we regress statistics that measure the reassortment intensity for a pair of segments of the flu genome on the body size of the host (a proxy for mean lifespan). Second, we infer phylogenetic networks (16) that summarize patterns of genetic ancestry (coalescences and reassortment) in a sample, and regress posterior mean reassortment rates on host body size. Finally, we discuss the consequences of viral recombination for emergence in a novel host.

## 2 Methods

We use a mechanistic mathematical model to characterize how the ecology of a reservoir host species affects the rate at which a focal pathogen (herein used interchangeably with virus) recombines. We then test predictions of the theory using publicly-available avian H5N1 sequence data. We describe each approach in detail below and provide a visual summary of the model and steps taken to analyze the sequence data in Figures 1 and 2, respectively.

**Figure 1:**
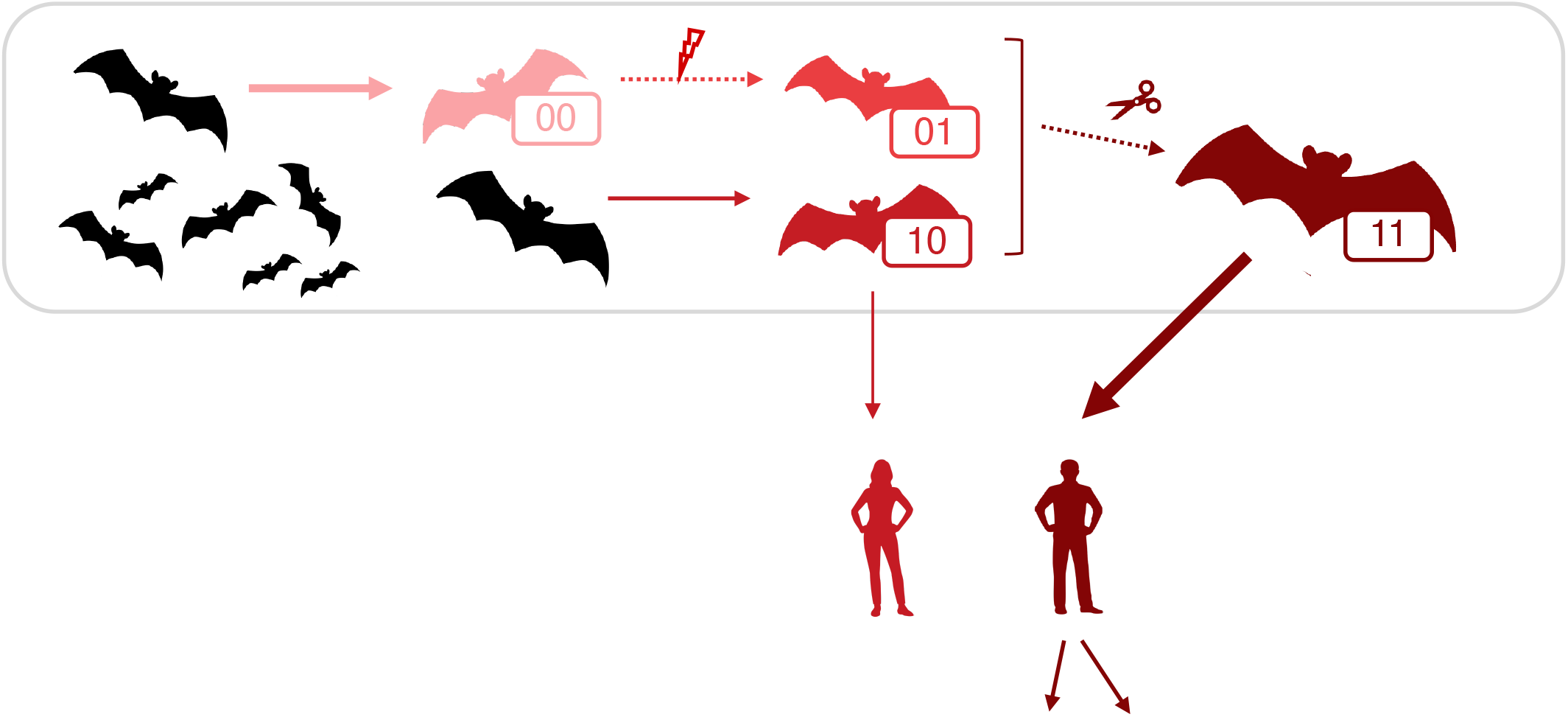
Model diagram. Susceptible individuals in the reservoir host population (in black) can be infected (shades of red) with one of the four possible pathogen genotypes (00, 01, 10, 11). Forward and back mutations occur at each locus and, when they fix, result in wholesale conversion of an infection to the mutant type. Recombination can occur when a host is infected with multiple genotypes and results in conversion of the infection to the recombinant type. The thickness of the solid lines corresponds to the intensity of transmission: the 00 genotype is most transmissible in the reservoir and least transmissible in humans, while the 11 genotype is most transmissible in humans but least transmissible in the reservoir.

**Figure 2:**
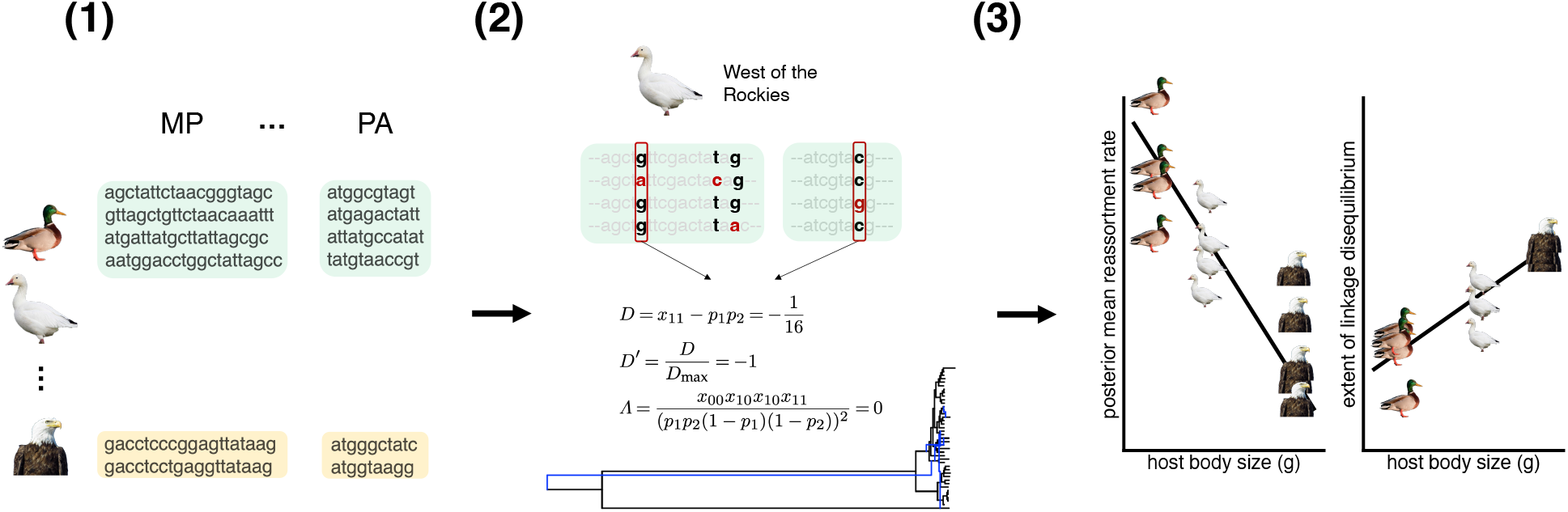
Analysis of publicly-available avian influenza sequence data. (1) Using GISAID, we retrieved all H5N1 sequences sampled in avian hosts in North America from December 2021-May 2024. We removed sequences without species-level host information, as well as those with nondescript sampling locations. (2) We grouped sequences by host species and “flyway”, and kept host-flyway combinations where *n* ≥ 50 to test model predictions for LD and where *n* ≥ 20 for reassortment. For each combination of host, flyway, and influenza genome segment (HA, NA, MP, PA, etc.), we sub-sampled and aligned the sequences using MAFFT (41). For all combinations of mutations found on different segments in a host and flyway, we calculated the population genetic statistics |*D*^*′*^| and Λ. An example calculation of these statistics is shown in the figure. Using CoalRe, we estimated reassortment networks and rates (per lineage per year). (3) To test predictions of the model, we regressed mean pairwise linkage disequilibrium measures and posterior mean reassortment rates on host body size. Predictions of our model are depicted in the figure.

### 2.1 Mathematical model

Our model extends the two-species, multi-strain Susceptible (S), Infected (I), and Recovered (R) framework used by Nuismer et al. (27) to understand how the propensity of a pathogen to spillover and emerge in a human population depends on the ecology of a host species that serves as a reservoir for the pathogen. The role that reservoir host traits play in shaping infectious disease dynamics, pathogen evolution, and disease emergence has been the focus of other modeling work (19; 28), but none of these studies considered the role of recombination in the emergence process. McLeod & Gandon (29) modeled recombination between co-infecting pathogen genotypes, but their goal was to clarify how the mode of action of an imperfect vaccine affects the joint evolution of virulence and vaccine escape. Our treatment of recombination in a co-infected host is similar, but the questions which we seek to answer are different. In particular, we use the model to characterize how the rate of viral recombination and the risk of disease emergence in humans differs between types of reservoir host. This is done, as in Nuismer et al. (27), by distinguishing host taxa based on mean lifespan, the rate of recovery from infection, and the rate of waning immunity. In the main text, we focus on lifespan, since among the three, this is most clearly a host trait and an empirical proxy is readily available. In the supplementary material, we explore how the rates of recovery and waning immunity (both likely to be an outcome of a combination of host and viral traits) affect the realized rate of recombination.

#### 2.1.1 Model assumptions and dynamics

In our model, individuals in the reservoir host population are tracked and distinguished based on their disease status and, among those which are infected at a given time, the genotype of the virus they carry. The density of susceptible reservoir hosts is denoted *S*. The density of individuals infected with genotype *g* is denoted *I*_*g*_. The density of hosts that have recovered from infection is denoted *R*. Throughout the paper, *I*_*T*_ = ∑_*g*_ *I*_*g*_ is used to denote the total density of infected hosts and *N* = *S* + *I*_*T*_ + *R* the reservoir population density.

We consider the simplest setting in which one can study patterns and consequences of viral recombination: when there are *n* = 2 loci underlying transmission in the reservoir host and humans. With two alleles (0 and 1) at each locus, four viral genotypes (*g* ∈ {00, 01, 10, 11}) must be tracked. The loci under consideration can represent single nucleotides, or longer stretches of non-recombining DNA or RNA. If loci fall on different segments, our treatment of recombination is equivalent to a model of reassortment. The alleles at each locus can represent specific combinations of mutations (or lack of such mutations) that affect the capacity of the virus to transmit in the reservoir and emerge in humans. We imagine these mutations affect phenotypes important in the cross-species transmission process, such as receptor binding and cell entry (30). As in Nuismer et al. (27), we assume there is a trade-off between transmission of the virus in the reservoir host and emergence in humans. In particular, the ‘1’ allele at each locus promotes transmission to and between humans but is costly for transmission in the reservoir host population.

In the model, coupled demographic, epidemiological, and evolutionary processes change the values of the state variables. As in Nuismer et al. (27), individuals are born at per-capita rate *b*, die at per-capita rate *d*, and experience density-dependent birth and death at rates *α* and *ρ*, respectively. When new individuals are born, they are born susceptible. Infection is modeled using mass action: susceptible individuals become infected with genotype *g* at rate *β*_*g*_*I*_*g*_. Individuals recover from infection, regardless of the infecting strain, at rate *γ*. Immunity is fully cross-reactive: once recovered, individuals cannot become infected with any genotype. Recovered individuals lose immunity at rate *δ*. Finally, there is no disease-induced mortality. This assumption permits straightforward comparisons with Nuismer et al. (27) and provides an approximation to a model with virulence when the reservoir species has co-evolved with the pathogen for an extended period of time (31).

During an infection, mutation and recombination can occur. The rates at which forward (0 → 1) and backward (1 → 0) mutation at locus *i* occur *and* fix within a host (resulting in conversion of the infection to the mutant type) are given by *µ*_*i*_ and *ν*_*i*_, respectively. Similarly, the rate at which recombinant genotype *ij* is created and fixes within a host is *σI*_*il*_*I*_*kj*_ (where *k*≠ *i, j*≠ *l*). The change in the density of individuals infected with genotype *g* = *ij* due to recombination is then given by *σ*_*ij*_ = *σI*_*il*_*I*_*kj*_ − *σI*_*ij*_*I*_*kl*_. The parameter *σ* is the rate infections with recombinant genotypes are produced (given infected individuals meet), and we assume that it does not depend on the particular makeup of a co-infection. This rate includes not only the rate recombination occurs in a co-infected individual, but the probability the recombinant type fixes within the host. The assumption that this rate does not depend on the makeup of the co-infection stems from the assumption that there are no differences in fitness at the within-host level, i.e., viral genotypes are exchangeable within each host. Although a simplification, this provides a tractable approximation to a more complex model that explicitly captures within-host dynamics. An explicit co-infection framework would be useful when investigating questions where the within-host dynamics have a major effect on the fate of mutant and recombinant genotypes as they spread between hosts.

The joint ecological, epidemiological, and evolutionary dynamics are described by the following system of ordinary differential equations for *S, I*_*ij*_, and *R*:

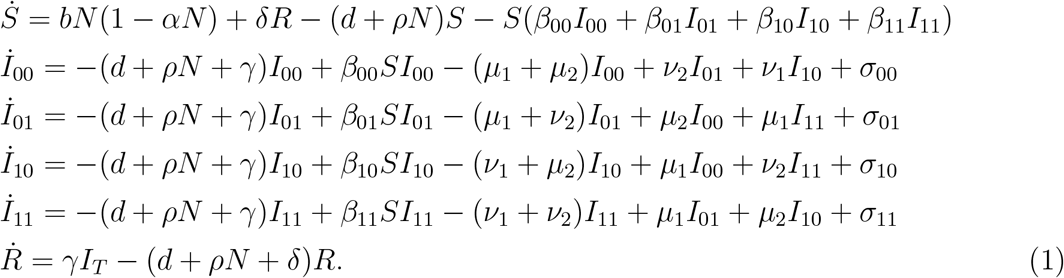

Assumptions regarding fitness of the viral genotypes in the reservoir host population are reflected in the parametrization of the transmission coefficients in the preceding system of equations. We assume the 00s transmit at a rate *β*_00_ and that carrying a 1 at locus *i* results in a multiplicative reduction in this rate by 1−*c*_*i*_. The double mutants experience a reduction in transmission by amount 1 − *c*_1_ − *c*_2_ + *e*, where *e* is the epistasis between mutations affecting transmission in the reservoir, i.e., departures in fitness from the additive action of alleles that arise from carrying 1s at both loci. (Note that this generalizes epistasis in the model of Nuismer et al. (27), where the assumption that alleles have independent multiplicative effects on transmission implies *e* = *c*_1_*c*_2_ *>* 0.) Positive values of *e* mean that the double mutant is more fit than would be expected under a model where the costs to transmission were additive; negative epistasis means that the double mutant is less fit than would be expected. The sign and magnitude of epistasis shape the extent to which associations between alleles are formed, i.e., how often ‘1’ alleles are found on the same genetic background relative to what is expected based on the allele frequencies. When epistasis does not reverse the sign of selection on the double mutant (i.e., *e* ≤ *c*_1_ + *c*_2_), as we assume here, transmission is greatest for the 00 genotype (i.e., *β*_00_ ≥ *β*_01_, *β*_10_, *β*_11_).

Parameterization of the transmission coefficients in terms of *c*_1_, *c*_2_, and *e* specifies the fitness landscape for the virus in the reservoir host population, but does not tell us anything about fitness of the different genotypes in humans. Our results regarding how host traits influence the rate of viral recombination are not sensitive to what the fitness landscape looks like in the human population, but in order to generate predictions regarding the role of recombination in emergence, we assume that there is a trade-off between transmission in the reservoir host and in humans. The reasons we assume a trade-off are two-fold. First, most mutations are deleterious. So, if mutations that are adaptive in another environment are drawn *randomly* from the distribution of fitness effects of new mutations, then they are most likely deleterious in the current host. Second, the assumption of a trade-off in transmission rates between host species has modest empirical support. There is evidence that destabilizing mutations, which are often deleterious, can facilitate future adaption by allowing receptor-binding proteins to recognize new hosts (30; 32). This is the justification of Nuismer et al. (27): if the optimal configuration for a protein involved in cell entry is different in humans than in the reservoir host, in which the virus may already be well-adapted, changes in the structure that enhance transmission to/between humans are likely detrimental to binding of the reservoir host receptor and, thus, transmission in the reservoir host.

#### 2.1.2 Evolutionary sub-system

The full model gives rise to an evolutionary sub-system that describes the frequencies of the four genotypes through time, *x*_*ij*_ = *I*_*ij*_*/I*_*T*_. Moreover, since these frequencies sum to one, it is enough to track *p*_1_ = 1 − *q*_1_ = *x*_11_ + *x*_10_, the frequency of 1 at the first locus; *p*_2_ = 1 − *q*_2_ = *x*_11_ + *x*_01_, the frequency of 1 at the second locus; and *D* = *x*_11_ − *p*_1_*p*_2_, the linkage disequilibrium (LD) between the alleles under consideration. LD measures how much more or less likely alleles at each locus are to be found on the same genetic background than expected by chance. *D* is the key variable of the evolutionary sub-model. It is through this quantity that recombination can influence the genotype frequencies. In fact, the terms involving recombination in the full model can be re-cast in terms of *D*:

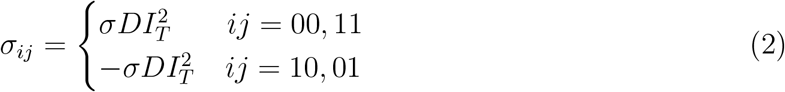

Thus, we need to analyze the behavior of LD to characterize feedbacks between host ecology and recombination and the consequences of recombination for disease emergence.

The evolutionary sub-model, which together with equations for the ecological variables *S, I*_*T*_ , and *R* is sufficient to characterize the dynamics of the full system, is

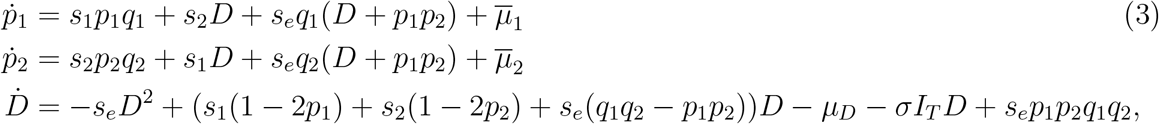

where 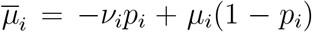 is the change in allele frequencies due to mutation, *µ*_*D*_ = −(*µ*_1_ + *µ*_2_ + *ν*_1_ + *ν*_2_)*D* is the change in *D* due to mutation, *s*_*i*_ = −*β*_00_*c*_*i*_*S* is the strength of selection at the *i*th locus, and *s*_*e*_ = (*β*_00_ + *β*_11_ − *β*_10_ − *β*_01_)*S* = *β*_00_*eS* is the strength of epistatic selection. Direct and epistatic selection increase with the number of susceptibles. This is because genotypes spread by successfully infecting susceptible hosts and differences in transmission rates are more important when there are more susceptibles to infect. A complete derivation of the equations can be found in Section S1.

#### 2.1.3 Comparing host populations and species

In order to quantify the degree to which hosts are short-vs long-lived, we define (following Nuismer et al. (27)) the per-capita rate of population turnover at equilibrium,

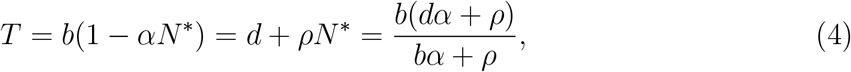

where the asterisks denote the value of a state variable at equilibrium. The mean lifespan of a host at equilibrium is 1*/T*. To quantify immune investment relative to a typical individual’s lifespan, we define Γ = *γ/T* and Δ = *δ/T* , i.e., the turnover-scaled rates of recovery and waning immunity, respectively. It is also useful to define the basic reproductive number,

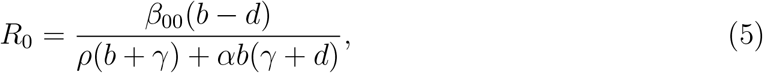

which defines the condition (*R*_0_ *<* 1) for stability of the disease-free equilibrium 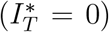 when there is no selection (i.e., viral genotypes are exchangeable).

Defining these quantities allows one to compare host populations/species with respect to measurable dimensions of their life history and immunology. However, care must be taken in deciding what parameters to hold constant. We focus on a simple, biologically-realistic scenario that allows for differences in lifespan (i.e., turnover) and equilibrium population densities (a function of turnover), while holding Γ, Δ, *β*_00_, and *R*_0_ constant. Holding Γ (resp., Δ) constant requires changing *γ* (resp., *δ*). Differences in equilibrium population density are, importantly, only possible because of density-dependence in birth and death rates; in the case where there is exactly no effect of population density on growth (*α* = *ρ* = 0) then equilibrium population sizes are determined by an initial condition and would have to be artificially varied with *T* to reflect the fact pace-of-life and population density co-vary (33; 34; 35). Here, a relationship between lifespan and population density arises naturally from the mechanistic description of density-dependent and -independent vital rates. As a result, changes to the unscaled parameters lead to changes in *R*_0_ unless *β*_00_ or one of the demographic parameters (*α, ρ, b, d*) is varied. Because there are more basal parameters than aggregates, there is flexibility in the choice of what to vary in order to keep all but one aggregate constant. All of our figures in the main text were made by changing *β*_00_ to keep *R*_0_ constant as *T* (i.e., turnover) is varied. And while the focus in the main text is on the effects of varying host lifespan, in the supplementary material we explore the effects of changing the scaled rates of recovery and waning immunity when host lifespan (i.e., turnover) and the transmission coefficent *β*_00_ and basic reproductive number *R*_0_ are constant.

### 2.2 Comparative genomics

Since our model makes predictions about how host traits (like mean lifespan) influence the rate of viral recombination and extent of LD, we sought to test predictions using available genome sequence data for highly pathogenic avian influenza (HPAI). The rationale for using avian influenza is that several conditions must be met to test the prediction for LD. (1) Genome sequences must be publicly-available and high-quality. (2) The pathogen must infect and sustain transmission in hosts that differ in mean lifespan. (3) The pathogen must undergo somewhat frequent recombination or reassortment. (4) The pathogen must be reasonably well-mixed at some spatial scale, so that it is possible to define closed populations.

HPAI is a system for which these conditions are satisfied. Complete genome sequences can be accessed readily through GISAID (36). Avian hosts differ in their life history, and most variation in mean lifespan is between rather than within species (37). The eight segments of the avian influenza genome (HA, NA, PB1, PB2, NS, NP, PA, MP) undergo frequent intra- and inter-subtype reassortment (38). Finally, previous work has shown that viral sequences are reasonably well-mixed within the four major flyways in North America (Pacific, Central, Mississippi, and Atlantic) for wild migratory birds (e.g., mallards, snow geese, bald eagles; (39)). Although there is structure within flyways, and somewhat frequent exchange of genotypes between flyways, the migration rate within flyways for avian influenza has — regardless of the sub-type considered — been shown to be higher than between flyways (38; 40). This makes avian influenza and, in particular the sub-type H5N1, a good candidate for testing our model predictions, in that it is possible to define populations based on i) the flyway in which a sequence was sampled and ii) the host species it was sampled in.

#### 2.2.1 Data processing and analysis

The steps we took to analyze the HPAI sequence data are described in Figure 2. Briefly, we used GISAID (36) to retrieve all H5N1 sequences sampled in avian hosts in North America from December 2021-May 2024. We removed sequences for which the host and/or location data were missing or not sufficiently detailed (i.e., at the state level or finer). After identifying the scientific names of each host species (and any aliases it has), we grouped sequences by host and if they were sampled East or West of the Rocky Mountains (i.e., by “flyway”). For analyses involving the calculation of LD, we kept host-flyway combinations where the number of sequences exceeded a sample size cutoff of *n* ≥ 50. For each host-flyway pair, we randomly sub-sampled genomes to avoid unequal bias in the estimates of LD statistics across host species. (This choice was made because there are more sequences for some host species, such as chickens and bald eagles, than others. Failing to sub-sample would result in unequal bias in LD statistics, making downstream comparisons difficult to perform and interpret.) For estimation of reassortment networks and rates using CoalRe, we used a sample size cutoff of *n* ≥ 20 and did not sub-sample. For each combination of host, flyway, and segment, we aligned the (possibly sub-sampled) sequences to each other using multiple sequence alignment software (MAFFT; (41)) using default parameters and then identified all polymorphic sites (excluding gaps and ambiguities).

For the LD analyses, we first identified occurrences of bi-allelic SNPs that were found on different segments and each at frequency *>* 5%. For all combinations of such SNPs, we computed two statistics quantifying the covariance between mutations on those segments:

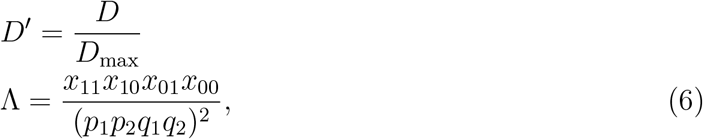

where *D*_max_ = min(*p*_1_*p*_2_, *q*_1_*q*_2_) when *D <* 0 and *D*_max_ = min(*p*_1_*q*_2_, *q*_1_*p*_2_) when *D >* 0. The first statistic, *D*^*′*^, measures departure from the free reassortment limit. It equals *±*1 when alleles are completely linked and 0 under free recombination (42). The normalization is necessary because the range of *D* depends on the allele frequencies, a known problem that makes comparisons of |*D*| across host taxa challenging (43). The second statistic, Λ, measures departure from the zero recombination limit and can be understood as a continuous analogue of the four-gamete test. It equals 0 when one of the genotypes is absent, 1 when recombination is free, and can assume values greater than 1 when there is strong positive epistasis (44). We then averaged these statistics over all retained combinations of SNPs and used linear regression to identify if body size, extracted from (45) as a proxy for the mean lifespan of a host, is associated with the mean value of each statistic.

After completing the LD analyses, we used CoalRe (16) to estimate reassortment rates and networks from the multiple sequence alignments associated to each combination of host, flyway, and influenza segment pair. These networks are graphical structures that summarize the patterns of genetic ancestry and reassortment in a sample of genomes; in population genetics, such networks are called ancestral recombination graphs. CoalRe uses Markov chain Monte Carlo to estimate parameters of the coalescent with reassortment. In each realization of the coalescent with reassortment (a network sampled in each MCMC iteration), lineages either find shared genetic ancestry (coalesce) backward in time or split (reassort) according to a constant-rate Poisson process. In addition to identifying the network structures (i.e., topologies and branch lengths) that are compatible with the genetic data, CoalRe provides estimates of various parameters that govern how sequence evolution unfolds on the branches (e.g., the rates mutations of various kinds arise and become fixed). Using the HKY+Γ4 nucleotide substitution model (46; 47) and a strict clock, we ran CoalRe on all combinations of host-flyway-segment pair for 200 million iterations. After all iterations were completed or the jobs timed out at 24 hours, we determined convergence (or lack of convergence) to the joint posterior distribution of parameters using effective sample sizes. We retained the combinations of host-flyway-segment pair runs where all parameters had estimated effective sample sizes *>* 200 (48) and removed the first 30 percent of posterior draws. Finally, we regressed posterior mean reassortment rates on host body size to determine if the reassortment rates needed to explain the genetic data conform to the predictions of the theory.

Sequence analyses were completed using a combination of custom bash and R scripts. Regressions and visualizations were completed in R (49).

## 3 Results

### 3.1 Recombination is most frequent and LD least extensive when hosts are short-lived, acutely infected, or lose immunity quickly

To understand how host traits such as mean lifespan affect recombination and the extent of LD, we used a series of approximations from theoretical population genetics. First, we used the “quasi-linkage equilibrium” (QLE) approximation, which assumes that recombination is stronger than selection and mutation, such that *D* settles into an equilibrium value much faster than the allele frequencies (50). Under this assumption (i.e., solving 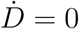 to leading order in a small quantity; assuming *σI*_*T*_ ≫ *s*_1_, *s*_2_, *s*_*e*_, *µ*_*i*_, *ν*_*i*_),

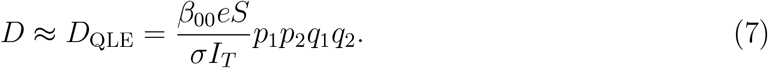

From this expression, it is clear that traits influence the equilibrium amount of LD in a couple of ways. First, differences in mean lifespan and the scaled rates of recovery and waning immunity can affect the equilibrium densities of susceptibles and infections. Changing the ecological variables affects both numerator and denominator in Eq. (7). But differences in the density of susceptibles affect the strength of selection on emergence-favouring alleles, *s*_*i*_ = − *β*_00_*c*_*i*_*S*, and the values allele frequencies take at equilibrium. How important those allele frequencies are in shaping the dependence of *D* on life history traits of the host, however, depends on the relative strengths of mutation and selection. Although all parts of the QLE expression are affected by host traits, further approximation reveals how *D* and recombination intensity (i.e., the denominator in (7)) vary with these parameters.

To develop more concrete predictions about the relationship between recombination, the extent of linkage disequilibrium, and host lifespan, we further assume that that selection is stronger than mutation. In Section S2 of the supplementary material, we show that, under the QLE and “strong selection, weak mutation” (SSWM) approximations,

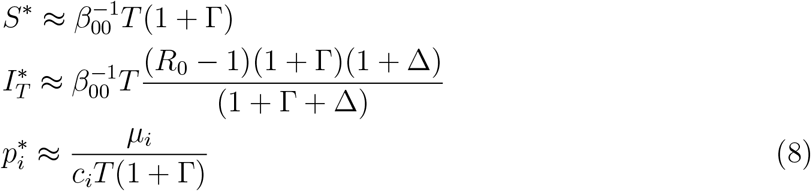

and, thus, the density of infections is increasing with turnover. This means that the rate of recombination (*σI*_*T*_) in viruses of hosts that are short-lived is greater than in hosts that are comparatively long-lived. Plugging in these expressions into Eq. 7,

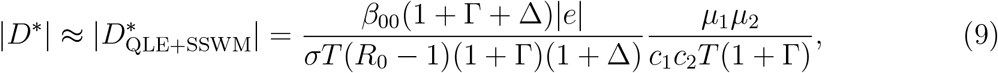

which is a decreasing function of turnover *T*. There are two reasons that increasing *T* decreases LD. First, increasing turnover increases the density of infections at equilibrium and, thus, the effective rate of recombination (Figures S1 and S2). Second, increasing turnover increases the density of susceptibles and, thus, the strength of direct selection on the 1 alleles, *s*_*i*_ = *β*_00_*c*_*i*_*S*^∗^. This is what Nuismer et al. (27) also found and, in fact, our expressions for the ecological equilibria match theirs under the QLE because LD is a second-order term. Because there are two alleles, increases in the magnitude of selection against the 1 alleles cancels the effect of increases in the extent of epitasis (also increasing with *S*^∗^). This gives rise to two terms in the previous expression which are decreasing with turnover, such that the magnitude of LD (in absolute value) is increasing with mean lifespan.

#### 3.1.1 QLE and SSWM approximations are robust

Having shown that the density of infections and rate of recombination decrease with mean lifespan and the extent of absolute LD increases with lifespan, we sought to determine if our results were robust to the assumptions used to develop these predictions. Based on numerical simulations of the full system (Eq. 1), it appears the predictions made under the QLE and SSWM approximations are qualitatively robust. Figure 3 (top panel) shows that absolute linkage disequilibrium is generally an increasing function of the mean lifespan of the host or, equivalently, a decreasing function of *T*. This is driven by reductions in the density of infections (Figure 3, bottom panel). Sensitivity analyses where parameter combinations were sampled using latin hypercube sampling show these results hold regardless of the relative strengths of mutation, recombination, and selection (Figures 3, S4, and S5).

**Figure 3:**
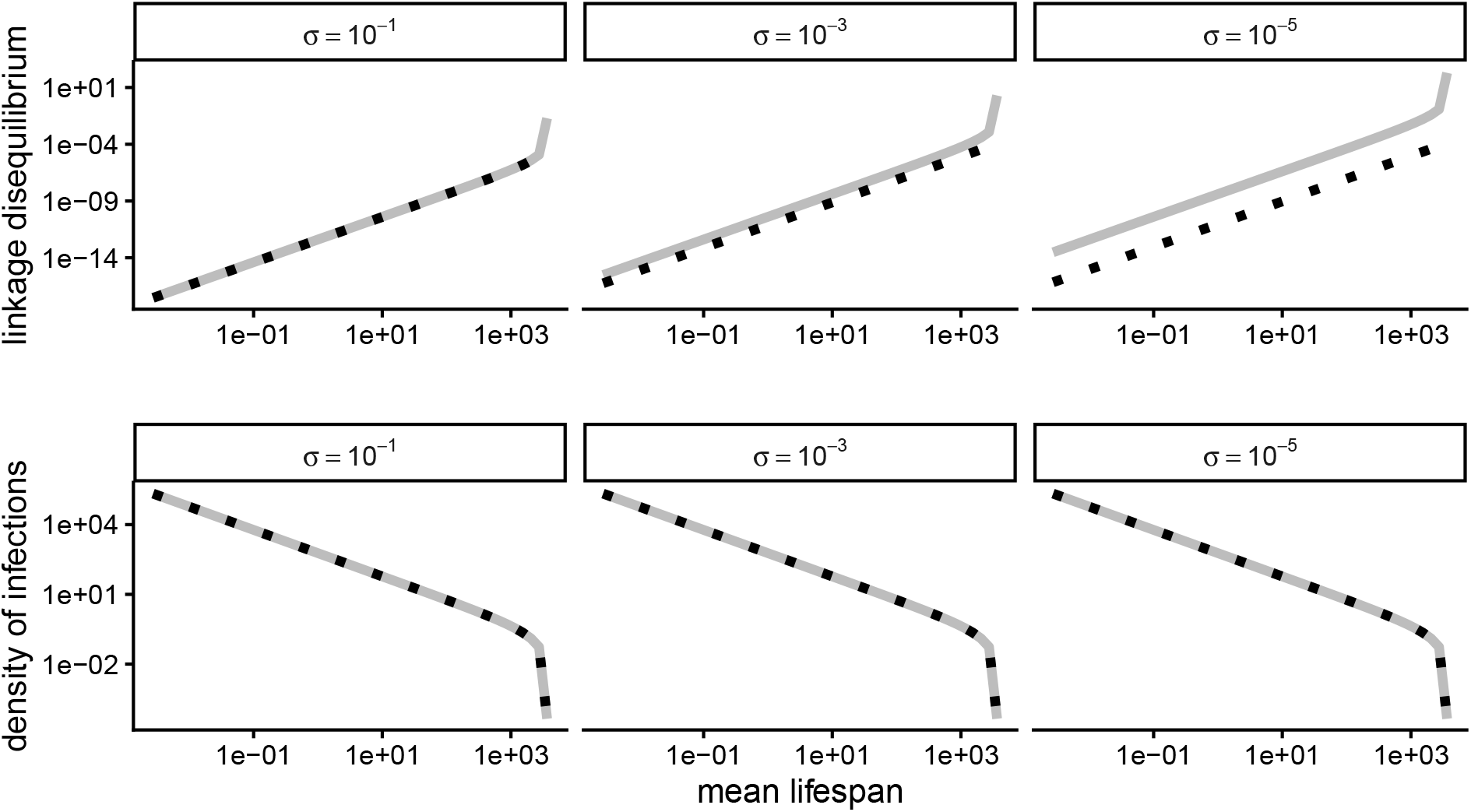
Robustness of predictions under the QLE and SSWM approximations. Solid, grey lines are the predicted relationship between host lifespan 1*/T* and equilibrium values of linkage disequilibrium *D* (equation 9; top panel) and infection density *I*_*T*_ (equation 8; bottom panel) for various recombination rates. Dotted, black lines are equilibrium values of *D* and *I*_*T*_ based on numerical solutions of 1. Parameters used: *R*_0_ = 5, Δ = 1, Γ = 25, *b* = 2*/*365, *d* = 1*/*3650, *ρ* = 0.0001, *c*_1_ = 0.2, *c*_2_ = 0.4, *e* = 0.5, *µ*_1_ = *µ*_2_ = 10^−5^, *ν*_1_ = *ν*_2_ = 10^−7^. Lifespan was varied through changes in *α. R*_0_, Γ, and Δ were held constant by changing *β*_00_, *γ*, and *δ*. Initial conditions used: *S*(0) = 0.99*N* ^∗^, *I*_00_(0) = 0.01*N* ^∗^, *I*_01_(0) = *I*_10_(0) = *I*_11_(0) = 0.

### 3.2 HPAI sequence data are consistent with model predictions

We used HPAI genome sequence data to test two model predictions: 1) recombination is greatest when hosts are short-lived and 2) linkage disequilibrium is an increasing function of mean host lifespan at equilibrium. Using body size as a proxy for mean host lifespan, we find that |*D*^*′*^| and Λ are associated with host body size as predicted (Figure 4; note that, because Λ measures departure from the zero-recombination limit, we predict it to be greatest when hosts are short lived). Although the effects have confidence intervals not overlapping zero, it is important to emphasize there is substantial residual variance that is not explained by host body size. Some of this variance could be due to variation in the acute-ness of infection, rate of waning immunity, or other host/viral traits for which we do not have measurements. Moreover, our predictions are about LD and related statistics, but these are composites of epistasis (*β*_00_*eS*) and recombination rate (*σI*_*T*_). It is not possible to de-convolute the effects of epistasis and recombination on patterns of linkage disequilibrium without, for example, making assumptions about the average epistasis for mutations on a segment pair.

**Figure 4:**
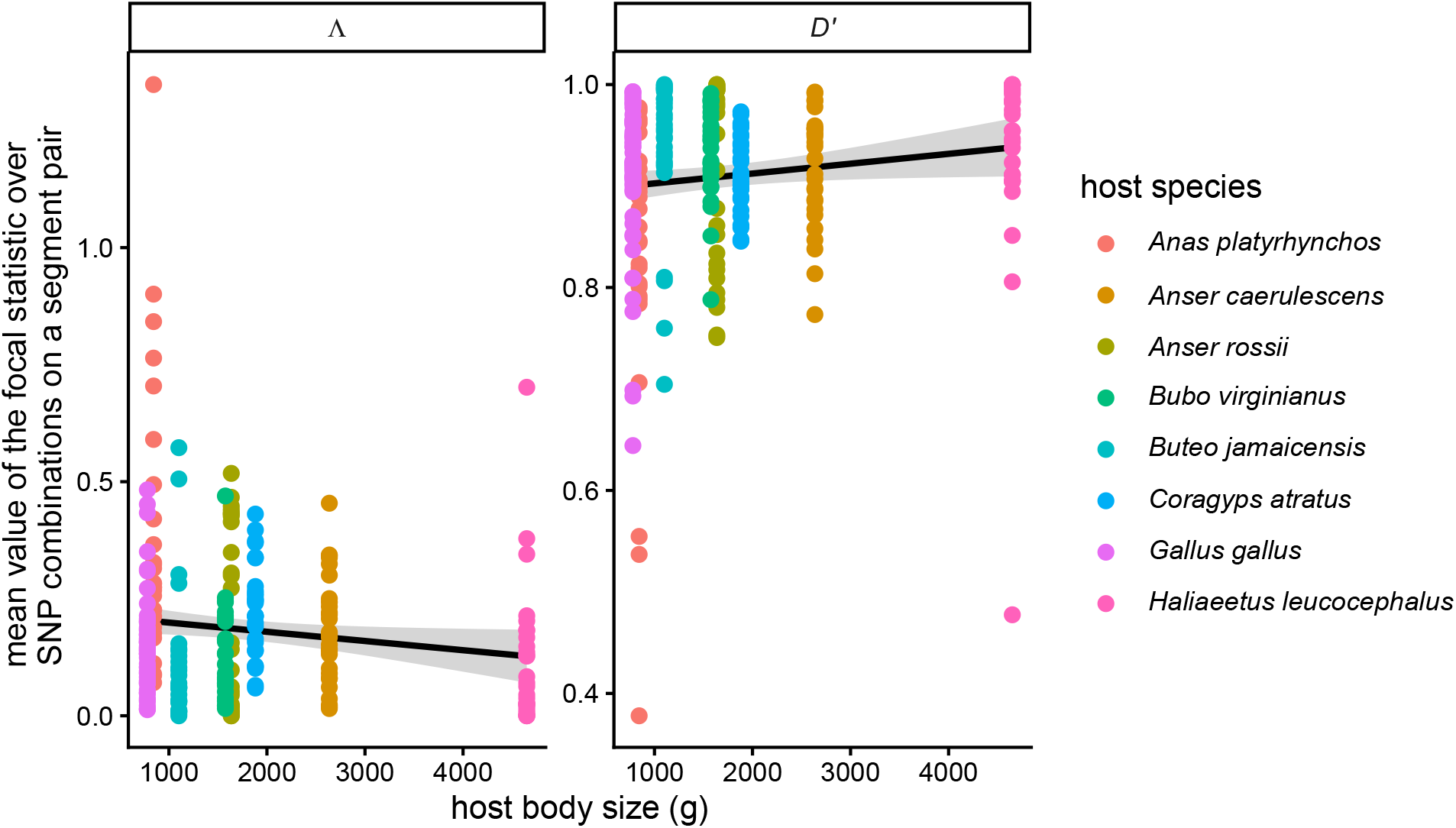
Sequence data are consistent with LD prediction: mutations on different segments of the avian influenza genome are, on average, more linked in long-lived (here, larger) host species. Each point corresponds to the mean Λ and *D*^*′*^ for a given segment pair in a host (color) and flyway. The estimated effect of size on |*D*^*′*^| was 9.7 *×* 10^−6^, with *p* = 0.0381. The estimated effect of size on Λ was −1.97 *×* 10^−5^, with *p* = 0.034.

Compelling evidence that |*D*^*′*^| and Λ are capturing signal in the data is that their values reflect our knowledge about avian influenza ecology and evolution. Namely, mutational combinations on the HA and the NA — which are believed to be prone to epistatic interaction due to their roles in cell entry and exit, respectively (51; 52) — have the highest mean |*D*^*′*^| and lowest mean Λ (over hosts) among all segment pairs (Figure S6). The fact that |*D*^*′*^| and Λ are extreme for mutations on the HA and NA is likely (and most parsimoniously) due to epistasis because, otherwise, it would have to be explained by differences in the reassortment rate between segment pairs. Second, the relatively large residuals of ducks (*Anas platyrhynchos*) compared to the regressions may be a signature of reassortment with other avian influenza sub-types. Since wild aquatic birds host a number of sub-types (53; 54), it is possible that (co-)infections with other subtypes and elevated inter-subtype reassortment explains the low values of *D*^*′*^ (resp., high values of Λ) observed in ducks (38).

Also in agreement with the predictions made by our model, the slope of the regression of posterior mean reassortment rate on body size was negative, with an estimated half as much reassortment per lineage per year in the largest hosts (roughly, the longest-lived) compared to the smallest (roughly, the shortest-lived). Visualization of the maximum clade credibility reassortment network reinforces that *per lineage* there are indeed far fewer reassortment events needed to explain the genetic data in populations of the largest bird (bald eagle, in our dataset) than the smallest bird (American crow, in our dataset; Figure 5).

**Figure 5:**
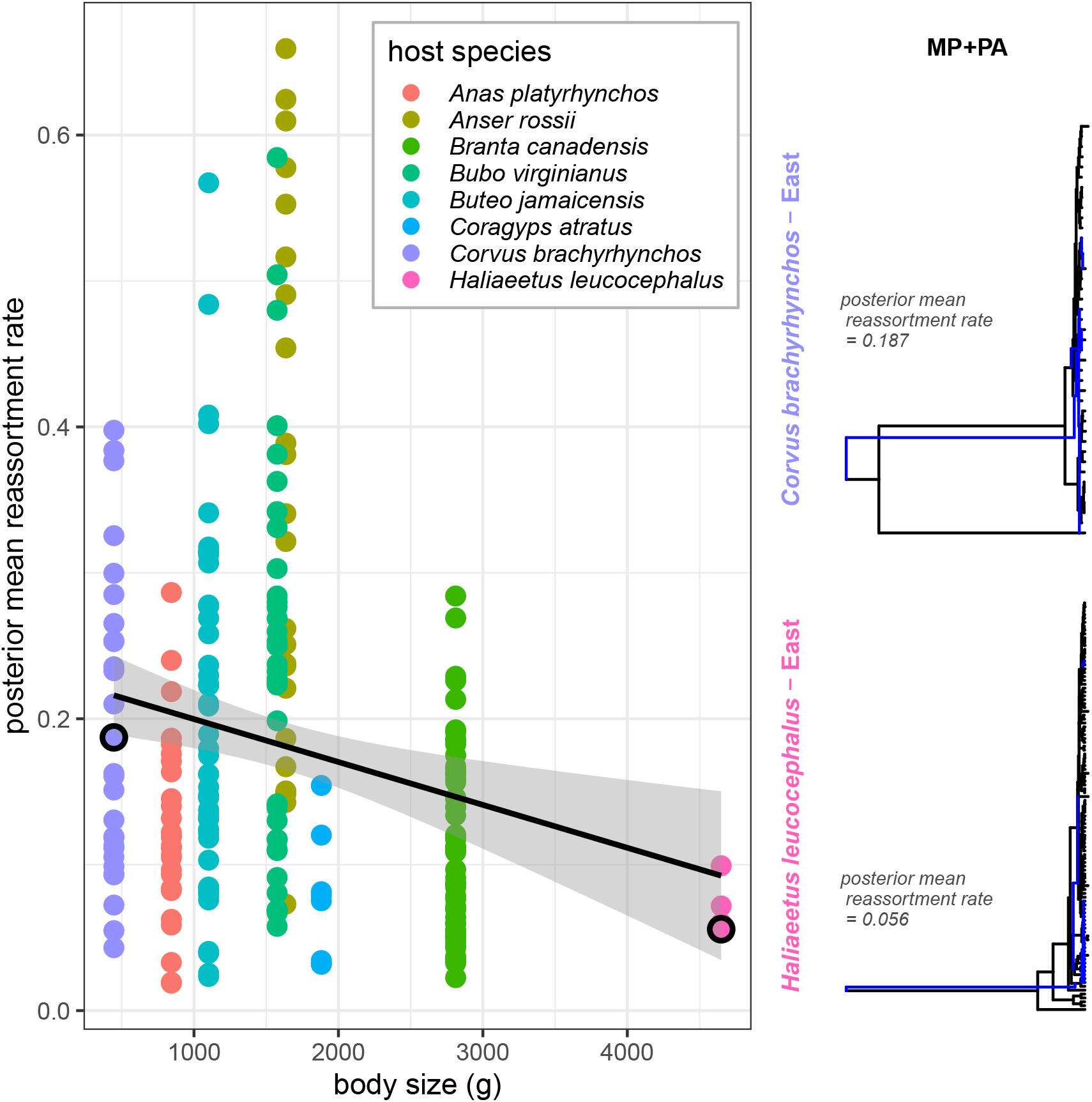
Sequence data are consistent with reassortment rate prediction: posterior mean reassortment rates estimated using CoalRe are highest in short-lived host species. In (A), each point corresponds to a combination of host species, flyway, and influenza genome segment pair where MCMC convergence diagnostics were satisfied, i.e., ESSs for all parameters were *>* 200. The estimated effect of body size on the posterior mean reassortment rate was − 2.9 *×* 10^−5^, with *p* = 0.00178. (B) shows the maximum clade credibility networks for a segment pair in the smallest- and largest-bodied hosts (i.e., MP+PA in bald eagles and American crow) visualized using the Reassortment Network Annotator (16) in the Bayesian Evolutionary Analysis Utility graphical user interface (BEAUti; default parameters) and R package ggret (55) Blue lines in the MCC reassortment networks are reassortants.

## 4 Discussion

Recombination in viruses is unique in that, due to coupling of recombination and co-infection (i.e., multiple strains simultaneously infecting the same host), its rate depends on the density of infected hosts (3). While co-occurrence of individuals in the same “patch” is required for sexual reproduction of any kind, co-infection is a stronger constraint on the rate at which recombination can take place. This is because an infected host is not just a patch. It exists in the context of other patches which are born, die, and turnover. Thus, the processes that influence number of patches (i.e., infected hosts) affect the frequency at which viral genetic material is recombined, which can in turn affect the number of patches. To explore how this feedback between host ecology and viral evolution (namely, recombination) plays out, we developed and analyzed a mathematical model. We found that the rate of viral recombination is greatest in hosts that are short-lived, all else equal. Intuitively, longer-lived hosts should have more opportunities to get secondary infections than shorter-lived hosts. We would therefore expect a virus population to have more opportunities for recombination in a longer-lived host population compared to a shorter-lived one. However, faster turnover of the shorter-lived population would give rise to a greater supply of susceptible individuals and potentially a higher infection rate, more secondary infections, and more opportunities for recombination. Our model results confirm this hypothesis. We also found that viruses of hosts that are acutely infected and which lose immunity quickly recombine more frequently than ones that are chronically infected and loose immunity slowly. We refer the reader to the supplementary material for those analyses and discussion of these findings.

### 4.0.1 Caveats

There are a number of limitations to both the model that we have built and the comparative genomic analyses that we have performed.

First, in the model, we ignore the complex and dynamic nature of immunity in many species by assuming viral genotypes are fully cross reactive (i.e., immunity to one gives rise to immunity to all). While this may not be a problematic assumption if, say, mutations affect cell entry, it is if they affect the way the virus navigates the immune system. Indeed, available evidence suggests that homosubtypic immunity is long-lasting (*>* 15 weeks in mallards (56)) but heterosubtypic immunity is not (i.e., re-infection with another subtype can occur within a few days (56; 57; 58)). With these data in mind, and in order to avoid violating the cross-reactivity assumption of the model, we only analyzed H5N1 data (i.e., sequences for the most prevent H5Nx subtype) in our comparative analyses.

Second, we make simplifying assumptions regarding virulence and within-host dynamics to arrive at the model we analyze (Eq. 1). One can imagine extending the model to include terms that correspond to disease-induced mortality (which is either genotype-independent or -dependent), or explicitly account for fitness differences in the replication of strains within hosts. Allowing for disease-induced mortality would influence mean lifespan at equilibrium, complicating fair comparisons across host taxa. Further, we expect predictions of the theory to be qualitatively unchanged *if* virulence does not depend on co-infection or the identity of infecting genotypes. This is because additional mortality in the infected class, if uniform, reduces the steady-state density of infections and results in greater magnitudes of LD than predicted in a model without virulence. Finally, we think our assumptions of avirulence and lack of polymorphism within hosts are reasonable to first order given that i) many zoonoses or potential zoonoses are co-evolved with their hosts (31) and ii) within-host dynamics tend to occur much faster than between-host dynamics (59; 60). That said, there is no question that the data we have chosen to use to test the prediction host lifespan is negatively associated with reassortment rate are in violation of the avirulence assumption. Indeed, HPAI is highly pathogenic in chickens and the 2.3.4.4b clade is thought to be more virulent in a number of hosts (61; 62). A promising avenue for future work is to explore to what extent virulence affects predictions for how host demography affects viral recombination.

Third, the data we have used to test predictions of the theory are the products of temporal and spatial structure, and frequent cross-species transmission. We have tried to control for structure by splitting sequences into different flyways and performing calculations for each host-flyway pair. However, this does not prevent the possibility that sequences in a host sampled west of the Rockies are the result of transmission from another host in the east. We imagine that some of the residual variance in the data is explained by the existence of structure and migration we could not account for and, for the LD data, the assumption that sequences are at equilibrium (i.e., ignoring heterogeneity in sampling dates).

### 4.0.2 What does this mean for emergence?

As recombination has been implicated in previous pandemics (17; 63; 64; 65), identifying the reservoir host taxa in which viral genotypes are likely to recombine is important. Our results suggest that sampling short-lived taxa could be a promising strategy to this end, given challenges measuring the magnitude of co-infection directly. Our results also show that the consequences recombination on emergence risk depend strongly on the sign of epistasis, the relative magnitudes of the allele frequencies and LD, and the genetic architecture of traits underlying cross-species transmission and emergence.

This risk can be quantified by calculating the “force of emergence” on humans, *λ*_emergence_ ∝ *I*_11_ = *x*_11_*I*_*T*_ , i.e., the rate at which emergent-capable genotypes establish in humans. Under the strong selection, weak mutation (SSWM) and quasi-linkage equilibrium (QLE) approximations that we have used to characterize the effects of host lifespan (as well as the duration of infection and rate of waning immunity in the supplementary material) on recombination rate and LD, the force of emergence is greatest when hosts are long-lived. Under these approximations, *p*_*i*_ = *O*(*ε*) and *D* ∼ *s*_*e*_*p*_1_*p*_2_ = *O*(*ε*). This means *λ*_emergence_ ∝ *I*_11_ = *x*_11_*I*_*T*_ = (*p*_1_*p*_2_ + *D*)*I*_*T*_ ≈ *p*_1_*p*_2_*I*_*T*_. Plugging in the approximate equilibria for the allele frequencies and density of infections, we have

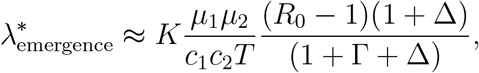

at equilibrium, where *K* is a constant of proportionality. Based on the expression, the force of emergence is decreasing with *T* , turnover. It is approximately independent of recombination rate because *D* ≪ *p*_1_*p*_2_ under the QLE and SSWM approximations; LD has already been broken down and breaking it down further has an inconsequential effect on the density of infections which are emergence-capable. The reason the equilibrium force of emergence is decreasing with turnover (increasing with mean lifespan) has to do with how the density of infections and the allele frequencies scale with *T*. Since increasing *T* increases the density of susceptibles and infections, it simultanously increases 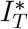 and decreases (*p*_1_*p*_2_)^∗^ by altering the strength of selection. Reduction in the frequency of 11s due to selection outweighs the increase in the density of all infections as *T* increases.

This analysis provides context for conclusions drawn by Nusimer et al. (27), who show with simulations that increasing the number of loci that underlie transmission in humans biases the force of emergence towards reservoir hosts that are longer-lived. The reason is that the probability of sampling a genotype with emergence-favouring alleles at all relevant loci is greatest when selection against 1s is weak, i.e., hosts are long-lived. With only one locus, the effects of ecology and evolution (favouring short- and long-lived hosts for emergence, respectively) cancel, such that the expression for the force of emergence does not depend on *T* , decreases with Γ, and increases with Δ. Our analysis of the two-locus model provides analytical justification for the observation that the force of emergence is increasing with host lifespan when more than one locus underlies emergence. This is because the QLE approximation we have used applies so long as the minimum recombination rate between a pair of loci is greater than the maximum selection coefficient for a emergence-favouring allele or epistatic coefficient for a pair of such alleles (66). Under the QLE,

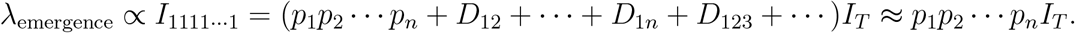

The sampling of 1s at each locus is roughly the product *p*_1_ · · · *p*_*n*_ due the fact that linkage disequilibria between mutations are lower-order terms. Assuming SSWM, the equilibrium allele frequencies are each inversely proportional to *T*. Then, given 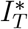 is proportional to *T* , the full force of emergence is decreasing with *T* when *n* ≥ 2. This approximation shows that, indeed, it becomes more difficult to sample 1s at all loci as reservoir hosts become shorter lived because selection against those alleles in the reservoir is stronger.

### 4.0.3 Outlook

Our results show that recombination need not be a prerequisite for emergence or even an especially important mechanism by which viruses evolve to sustain transmission in new hosts. Since *D* is a second-order term when recombination is stronger than selection and mutation, varying the recombination rate has little effect on the density of emergence-capable genotypes. This is in agreement with the verbal arguments in Simon-Loriere & Holmes’s review of RNA virus recombination (2). Our findings are also in agreement with Geoghegan et al. (67), who found that recombination/re-assortment frequency did not predict emergence risk compared to other features (e.g., if the virus was enveloped or not). Interestingly, Geoghegan et al. (67) also found that viruses that cause chronic infections are more likely to transmit successfully between humans, in line with the prediction for the force of emergence under the QLE and SSWM approximations (as discussed in Section 4.0.2).

How do we reconcile the observation that several pandemics have been facilitated by recombination and reassortment with the fact that, often, the effects of recombination on the force of emergence are second-order in nature? One possibility is that recombination shortens the waiting time to the first successful emergent-capable strain when intermediate genotypes are unable to emerge themselves and are at relatively low frequencies (68). This is a possibility which is not considered in our deterministic model of allele frequency and LD dynamics, but would be interesting to explore. Recombination could also play an important role in emergence when both beneficial and deleterious variants circulating, or when there is sign epistasis (i.e., genotypes must cross fitness valleys (69)). Finally, it is possible that recombinant and reassortant genotypes possess novel antigenic properties in the reservoir host and can spread to high frequencies when immunity is not fully cross-reactive; such a setup would likely give rise to more complex kinds of LD dynamics and consequences of recombination on viral evolution and infection dynamics (70). Mathematical modeling of reservoir ecology, viral evolution, and disease emergence that i) integrates the stochastic production of double mutants via mutation and recombination, ii) relaxes assumptions regarding immunity and infection, and iii) explores the effects of recombination under realistic distributions of fitness effects of new mutations in the reservoir host and humans (71) would help clarify when recombination matters for emergence.

Nevertheless, the growing recognition that recombination varies dramatically across the Tree of Life (72) suggests it is an exciting time to study how recombination varies in viruses. Indeed, a growing number of studies explore this variation (in rate, localization on the genome, etc.) in RNA viruses (10; 73). We too explore the causes and consequences of this variation and find, like Romero & Feder (10), that there are predictable signatures of density-dependence in the process that are left in present-day genomes.

## Ethics

This work did not require ethical approval.

## Data accessibility

Scripts used to complete data analysis and visualization can be found in this GitHub repository: https://github.com/metekaanyuksel/host-constraints-viral-rec.

## Declaration of AI use

We have used AI-assisted technologies to aid in scripting, including plotting.

## Acknowledgments

We gratefully acknowledge all data contributors, i.e., the Authors and their Originating laboratories responsible for obtaining the specimens, and their Submitting laboratories for generating the genetic sequence and metadata and sharing via the GISAID Initiative, on which this research is based. Computations were performed on the Trillium supercomputer at the SciNet HPC Consortium. SciNet is funded by Innovation, Science and Economic Development Canada; the Digital Research Alliance of Canada; the Ontario Research Fund: Research Excellence; and the University of Toronto.

We thank Else Mikkelsen and members of the Osmond and Mideo labs for helpful conversations on the subject of this work. We thank Louise Moncla for help accessing the HPAI sequence data, Nicola Muller for advice for using CoalRe, and Maria Maltepes for useful discussions about reassortment network inference. Finally, we thank reviewers of this work for their careful reading of the text and suggestions.

## Supplementary material

### S1 The model

To understand how host ecology affects the intensity of viral recombination and emergence, we develop and analyze a mathematical model that couples host ecology and viral evolution (selection, mutation, and recombination). The notation and assumptions are described in the main text. The density of susceptible hosts, *S*, hosts infected with genotype *ij, I*_*ij*_, and recovered individuals *R* are governed by the following system of ODEs:

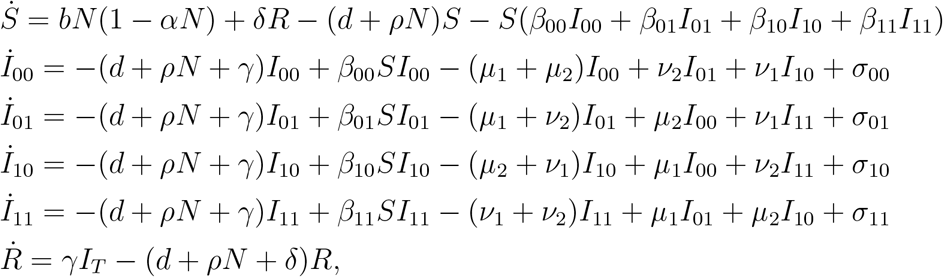

where *σ*_*ij*_ = *σI*_*il*_*I*_*kj*_ − *σI*_*ij*_*I*_*kl*_ (*k*≠ *i, j*≠ *l*).

#### S1.1 Derivation of evolutionary sub-system

Let *x*_*ij*_ = *I*_*ij*_*/I*_*T*_ be the frequency of the *ij* genotype among all infections. Then

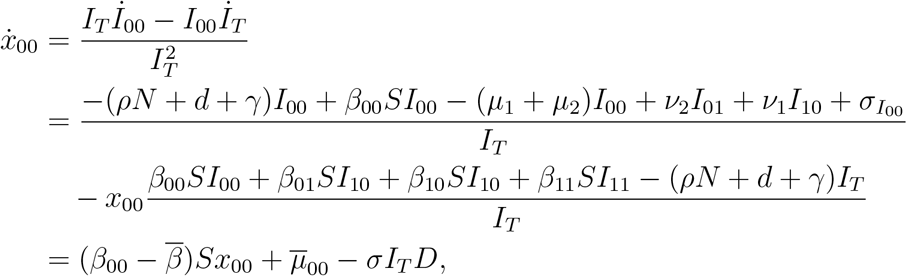

where 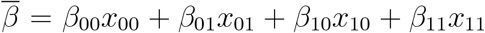 is the mean transmission rate in the reservoir and 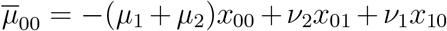 is mean change in the frequency of 00 infections due to mutation. The equation shows that the relative (dis)advantage in transmission 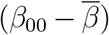, rates of recurrent mutation 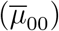, sign and magnitude of LD, and total number of infected individuals sets the change in frequency of the 00 genotype in the reservoir.

The full system of genotype frequencies is

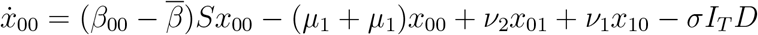

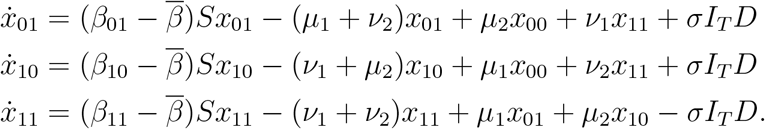

Now, let *p*_1_ = 1 − *q*_1_ = *x*_11_ + *x*_10_ be the frequency of 1s at the first locus and *p*_2_ = 1 − *q*_2_ = *x*_11_ + *x*_01_ be the frequency of 1s at the second locus. The system describing how genotype frequencies change can then be recast in terms of *p*_1_, *p*_2_ and *D* = *x*_11_ − *p*_1_*p*_2_.

In particular,

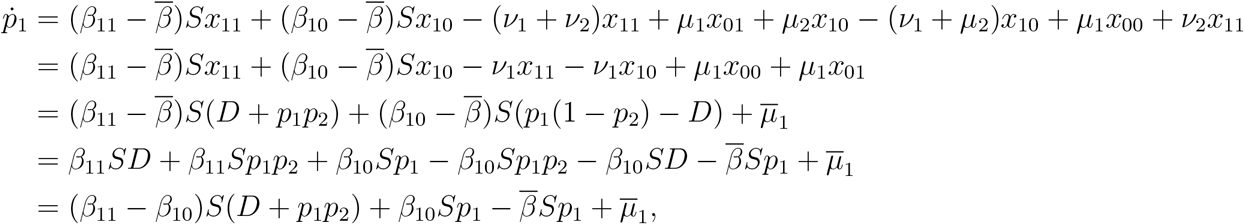

where 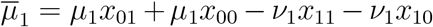 is the average change in frequency of 1s at the first locus due to mutation. Expressing the genotype frequencies in terms of *D, p*_1_, *p*_2_,

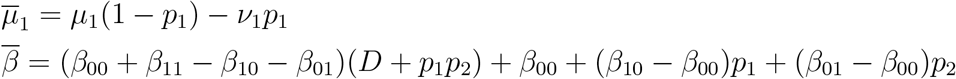

Ignoring mutation and recombination, the per-capita growth rate of *ij* infections is

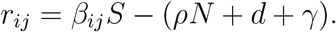

Following McLeod & Gandon (29), it is useful to define *s*_1_ = *r*_10_ − *r*_00_ = (*β*_10_ − *β*_00_)*S* and *s*_2_ = *r*_01_ − *r*_00_ = (*β*_01_ − *β*_00_)*S*, the selection coefficients for alleles of interest (here, 1s) at each locus. Since the number of susceptibles changes as individuals acquire infections of each type, lose immunity, and demographic changes unfold, the selection coefficients are time-varying. It is also useful to define epistasis in fitness, *s*_*e*_ = *r*_11_ +*r*_00_ − *r*_10_ − *r*_01_ = (*β*_00_ +*β*_11_ − *β*_10_ − *β*_01_)*S*. Ignoring mutation,

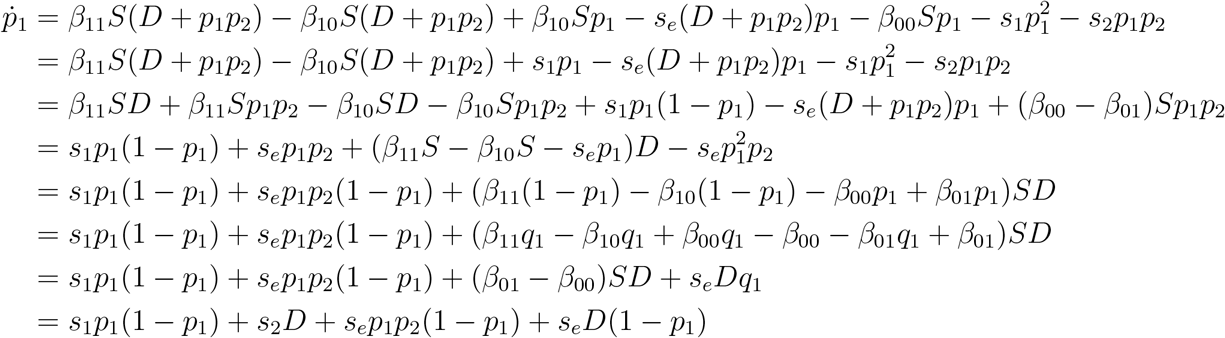

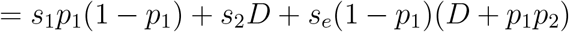

Similarly,

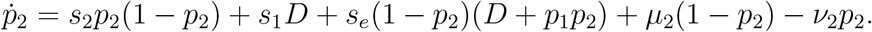

The first and second terms correspond to direct and indirect (i.e., linked) selection on each locus, respectively. The third terms describe change in *p*_*i*_ due to epistasis.

Moreover,

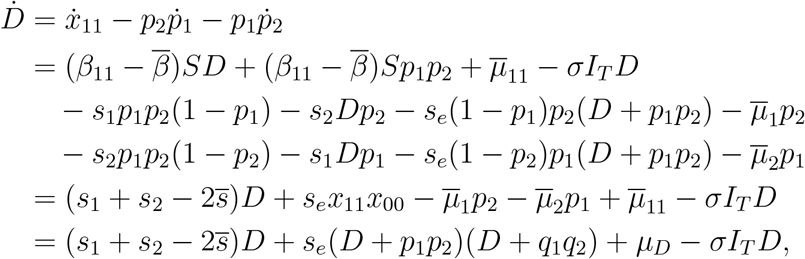

where 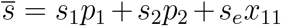 and 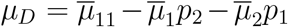. The change in LD due to mutation can be expressed in terms of the genotype and allele frequencies as follows:

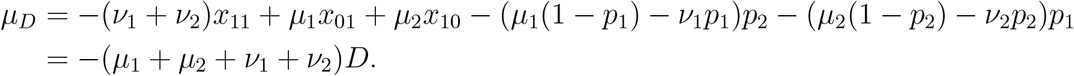

The full set of evolutionary equations, expressed in terms of allele frequencies and the linkage disequilibrium, is thus

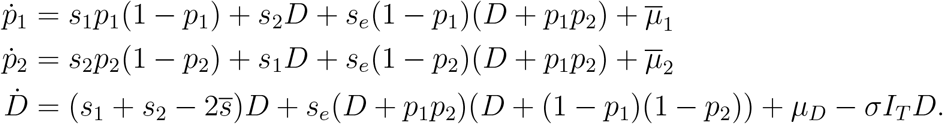

### S2 The behavior of LD at equilibrium

As described in the main text, we predict how recombination and LD scale with host traits using the quasi-linkage equilibrium (QLE) and strong selection, weak mutation (SSWM) approximations. If recombination is stronger than selection and mutation, i.e., *σI*_*T*_ ≫ *s*_1_, *s*_2_, *s*_*e*_, *µ*_*i*_, *ν*_*i*_, then LD changes on a fast timescale (relative to the allele frequencies) and can be assumed to be at a quasi-steady state *D*_QLE_. Under the QLE, linkage is small enough that it can be ignored when solving for the allele frequencies and, in our case, the joint eco-evolutionary equilibrium. Setting 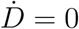 and rearranging terms,

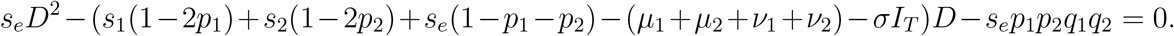

Treating mutation, selection, and epistasis as *O*(*ε*), expanding *D* = *D*_1_*ε* + *D*_2_*ε*^2^ + · · · + *O*(*ε*^*n*^) in a perturbation series in *ε*, and solving to leading order, one has

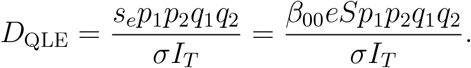

Making the additional assumption that selection is strong compared to mutation, *D* ≪ *p*_1_, *p*_2_ = *O*(*ε*) means 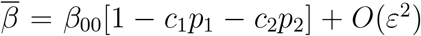. The approximate equilibrium allele frequencies are found by solving 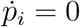 to first order, and are given by

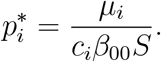

Plugging this into the equations for the ecological variables, we see that the approximate equilibrium values of *S, I*_*T*_ , and *R* satisfy

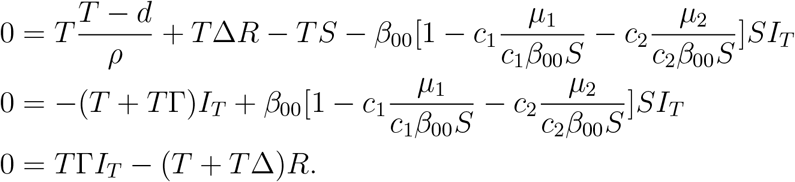

Thus,

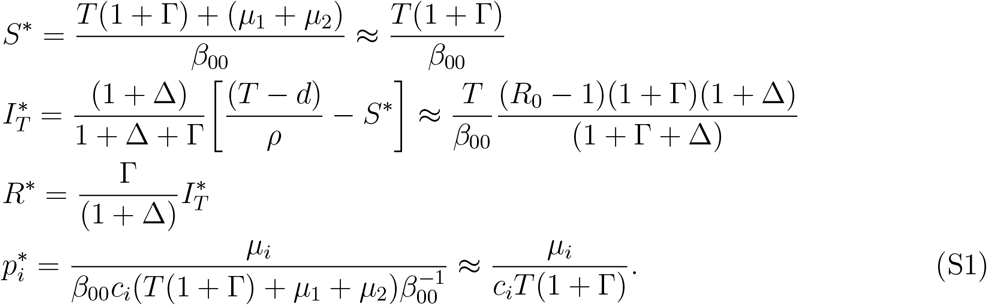

So

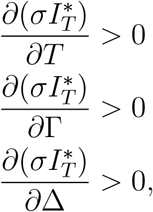

i.e., the effective rate of recombination at equilibrium is increasing with turnover (decreasing with host lifespan), the scaled rate of recovery, and the scaled rate of waning immunity.

Plugging in Eq. S1 into the preceding expression for *D*_QLE_,

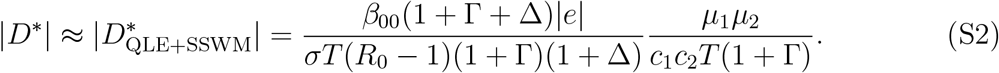

Thus,

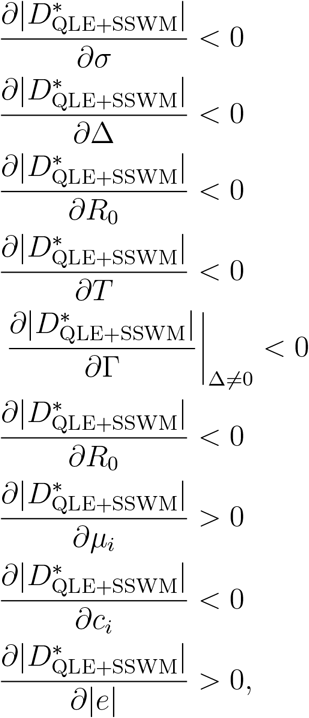

assuming that every other term in Eq. S2 is constant as *T* , Γ, or *R*_0_ are varied. See Figures S1-S5 for the relationships where *β*_00_ changes with *T* , Γ, or *R*_0_. Note that the expression for the ecological variables and allele frequencies under the QLE and SSWM match the equilibria in the “ecology-only” and single-locus models of Nuismer et al. (27).

The density of infections at equilibrium is increasing with Γ (and hence the effective rate of recombination is increasing and LD is decreasing) because, to keep *R*_0_ and the other aggregate variables constant in the comparison, the basal transmission rate *β*_00_ must increase. The greater basal transmission rate in host populations where individuals recover faster gives rise to a greater density of infections. Similarly, the density of infections at equilibrium is increasing with Δ because faster loss of immunity replenishes the pool of susceptible individuals and thus increases the rate new infections are generated.

**Figure S1:**
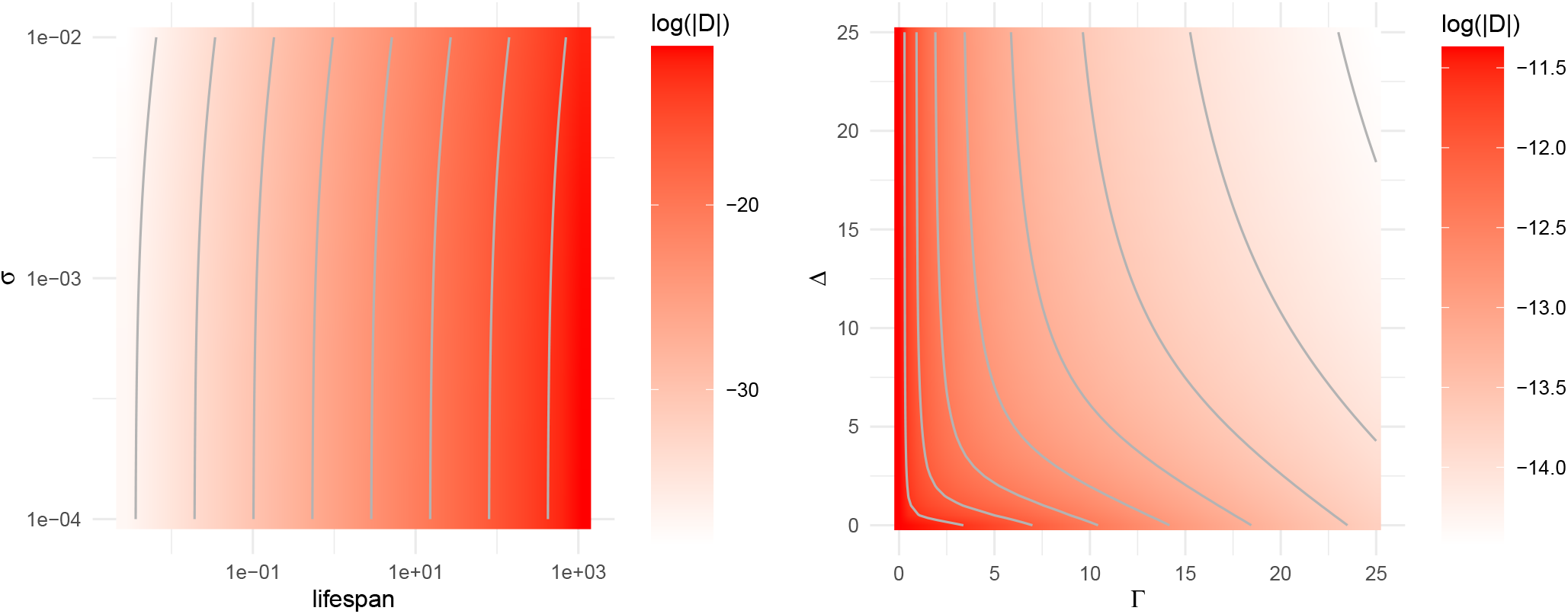
Under the QLE and SSWM approximations, |*D*^∗^| is an increasing function of host lifespan 1*/T* and a decreasing function of the scaled recovery rate Γ, the scaled rate of waning immunity Δ, and effective rate of recombination *σ*. Parameters used except for when varied: *σ* = 10^−3^, *R*_0_ = 5, Δ = 1, Γ = 25, *T* = 1*/*365, *b* = 2*/*365, *d* = 1*/*3650, *ρ* = 0.0001, *c*_1_ = 0.2, *c*_2_ = 0.4, *e* = 0.5, *µ*_1_ = *µ*_2_ = 10^−5^, *ν*_1_ = *ν*_2_ = 10^−7^. Initial condition: *S*(0) = 0.99*N* ^∗^, *I*_00_(0) = 0.01*N* ^∗^, *I*_01_(0) = *I*_10_(0) = *I*_11_(0) = 0.

**Figure S2:**
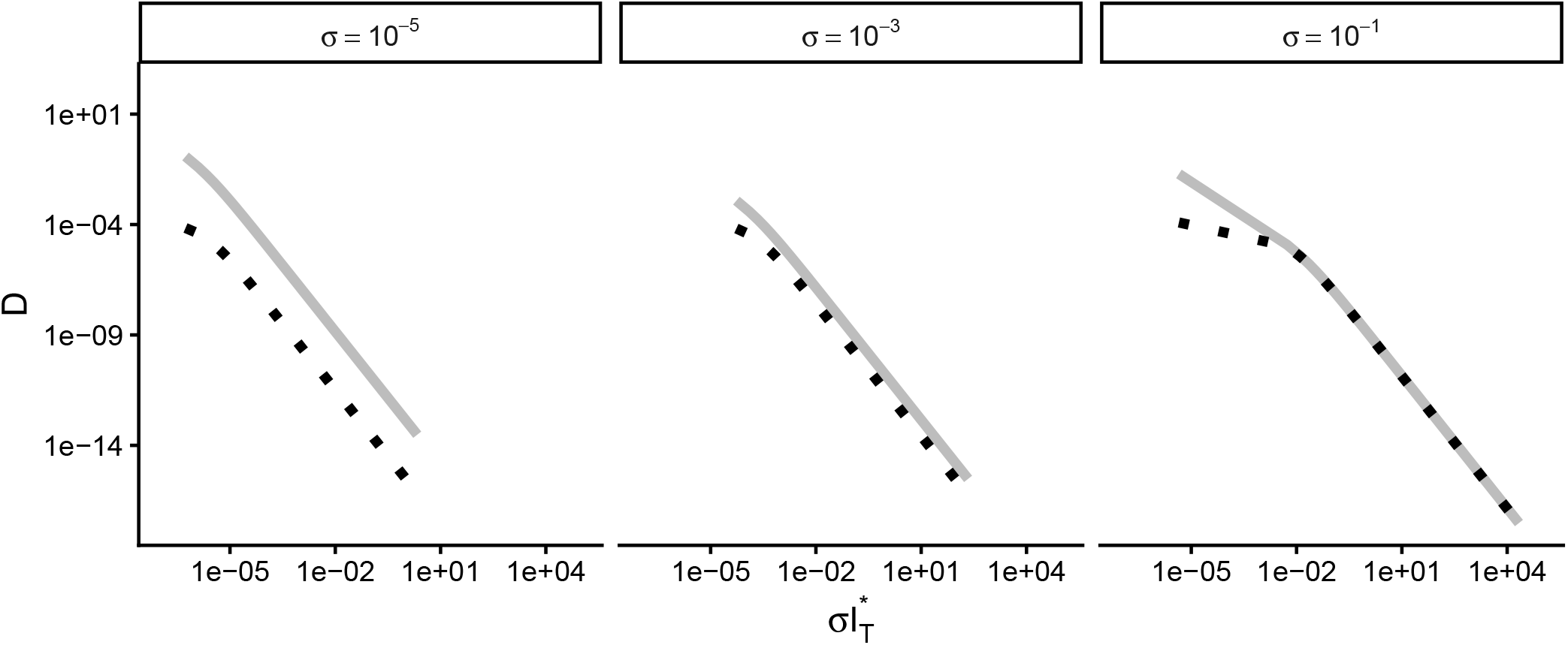
Relationship between *D*^∗^ and 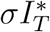 for various values of lifespan 1*/T* (which changes 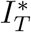) and the effective recombination rate *σ* (facets). Dotted lines correspond to equilibrium values from numerical simulations. Solid lines correspond to equilibrium values of *D* and *σI*_*T*_ under the QLE and SSWM approximations (Equations S2 and S1). The approximation does well with large values of *σ*. When *σ* is small, the approximation still captures the qualitative behavior. Parameters used: *R*_0_ = 5, Δ = 1, Γ = 25, *b* = 2*/*365, *d* = 1*/*3650, *ρ* = 0.0001, *c*_1_ = 0.2, *c*_2_ = 0.4, *e* = 0.5, *µ*_1_ = *µ*_2_ = 10^−5^, *ν*_1_ = *ν*_2_ = 10^−7^. Initial condition: *S*(0) = 0.99*N* ^∗^, *I*_00_(0) = 0.01*N* ^∗^, *I*_01_(0) = *I*_10_(0) = *I*_11_(0) = 0.

**Figure S3:**
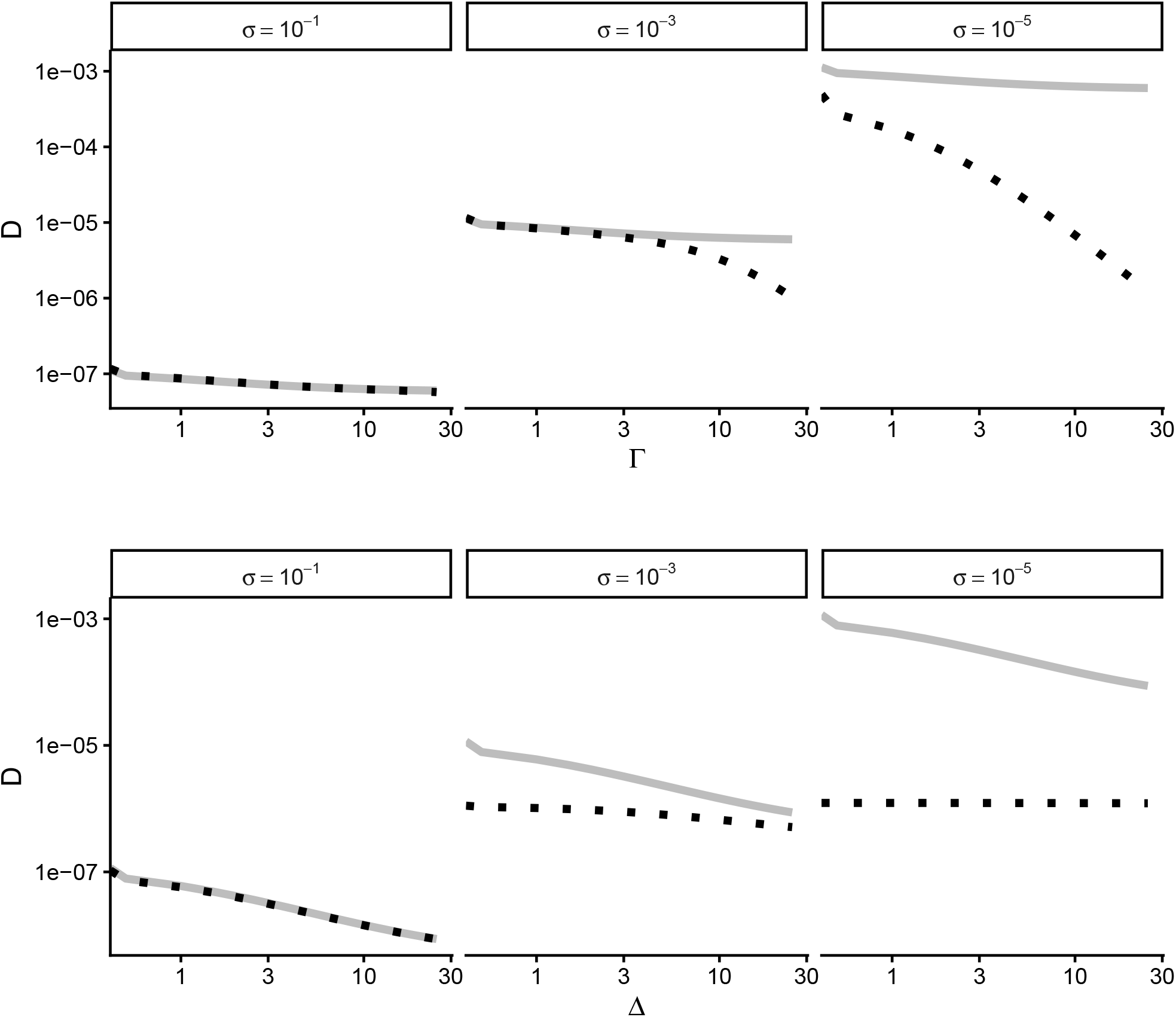
Robustness of predictions for *D* as a function of Γ and Δ under the QLE and SSWM approximations (equation S2) Solid lines are the predicted (approximate) relationship between the rates of waning immunity and recovery and equilibrium LD under various effective recombination rates (*σ* = 10^−6^, 10^−4^, 10^−2^). Dotted lines are equilibrium values of *D* based on numerical solutions. Parameters used except for when varied: *R*_0_ = 5, Δ = 1, Γ = 25, *b* = 2*/*365, *d* = 1*/*3650, *ρ* = 0.0001, *c*_1_ = 0.2, *c*_2_ = 0.4, *e* = 0.5, *µ*_1_ = *µ*_2_ = 10^−5^, *ν*_1_ = *ν*_2_ = 10^−7^. Initial condition: *S*(0) = 0.99*N* ^∗^, *I*_00_(0) = 0.01*N* ^∗^, *I*_01_(0) = *I*_10_(0) = *I*_11_(0) = 0.

### S3 Sensitivity analysis

To assess the sensitivity of the prediction that equilibrium values of |*D*| are increasing with host lifespan, we used Latin hypercube sampling (LHS; (74)) to explore regions of parameter space where the assumptions of the QLE and SSWM approximations break down.

**Table S1:** Range of parameter values used in sensitivity analysis. *α* was determined based on the equilibrium population size *N*^∗^ = (*T* − *d*)/*ρ* for each value of host lifespan used.

| Parameter | Description | Range |
| --- | --- | --- |
| $R_0$ | Basic reproductive number | [0, 20] |
| $\Delta$ | Turnover-scaled rate of waning immunity | [0, 50] |
| $\Gamma$ | Turnover-scaled rate of recovery | [0, 50] |
| $b$ | Per-capita birth rate, density-independent | [0, 10/365] |
| $d$ | Per-capita death rate, density-independent | [0, 1/3650] |
| $\rho$ | Per-capita death rate, density-dependent | $[10^{-5}, 10^{-3}]$ |
| $c_1$ | Cost of allele 1 at locus 1 on transmission in the reservoir | [0, 0.99] |
| $c_2$ | Cost of allele 1 at locus 2 on transmission in the reservoir | [0, 0.99] |
| $e$ | Epistasis between 1 alleles on transmission in the reservoir | $[-0.99, 0.99]$ |
| $\mu_1$ | $0 \rightarrow 1$ mutation rate at locus 1 | $[0, 10^{-2}]$ |
| $\mu_2$ | $0 \rightarrow 1$ mutation rate at locus 2 | $[0, 10^{-2}]$ |
| $\nu_1$ | $1 \rightarrow 0$ mutation rate at locus 1 | $[0, 10^{-2}]$ |
| $\nu_2$ | $1 \rightarrow 0$ mutation rate at locus 2 | $[0, 10^{-2}]$ |
| $\sigma$ | Effective rate of recombination, given infected individuals meet | [0, 1] |

Using the lhs package in R (75; 76), we randomly generated 1000 combinations of all parameters except host lifespan. Ranges for the parameters varied are described in Table S1. For each combination of parameters where *R*_0_ *>* 1, 1 − *c*1 − *c*2 + *e >* 0, and *e* < *c*_1_ + *c*_2_, we solved the multi-strain SIRS model (Eq. 1) to equilibrium for host lifespans from 1 day to 10 years, using the ode function in the R package ‘deSolve’ (77). Equilibrium values were those at the end of a 100 year simulation. Simulations where the steady-state value of |*D*| was less than 10^−16^ or the solver was unable to obtain the solution to the system at 100 years (= 36500 days) were discarded. The log-transformed equilibrium values of absolute linkage disequilibrium were then regressed on log-transformed values of lifespan. In Figure S4, estimates of the effect of log lifespan on log absolute LD are plotted (with confidence intervals) on the *y* axis and the associated *p*-values on the *x* axis. Values above the red line correspond to a relationship in the direction predicted under the QLE and SSWM approximations. Values to the left of the blue line have *p <* 0.05. All estimates of the effect of lifespan on the equilibrium magnitude of LD are in agreement with the prediction made under the QLE and SSWM approximations, i.e., that |*D*| is an increasing function of host lifespan (decreasing with turnover). This can also be seen in Figure S5, where the relationships between log-transformed values of equilibrium |*D*| and lifespan at each of the combinations of parameter values generated using LHS are plotted.

**Figure S4:**
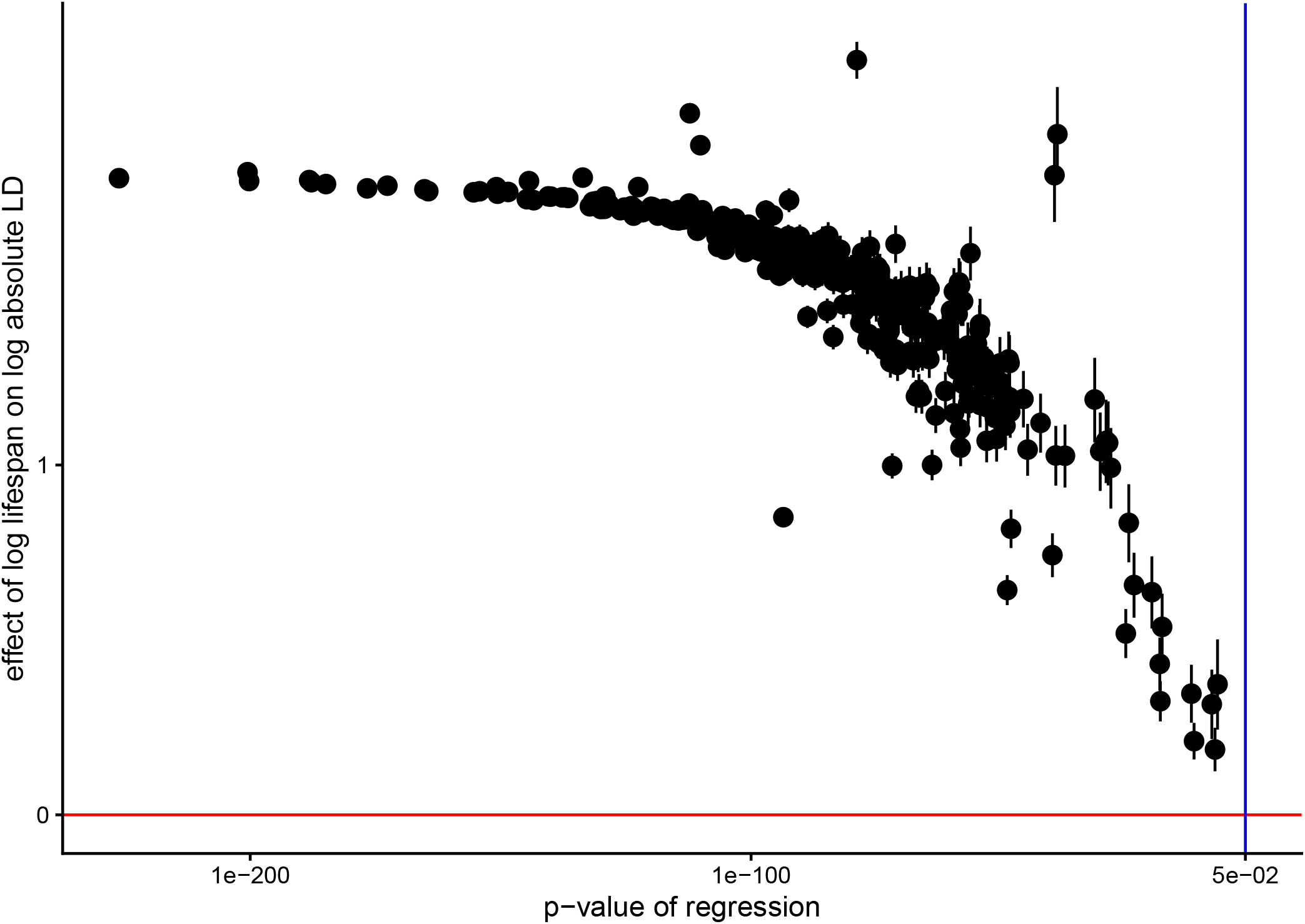
Robustness of the prediction that |*D*| at equilibrium increases with mean lifespan of a host to the assumptions of the QLE and SSWM approximations. As described in Section S3, Latin hypercube sampling was used to generate random combinations of parameters. Host lifespan was then varied and simulations at each combination of parameters carried out. Log absolute LD at equilibrium were regressed on log lifespan. Estimates of the effect size are plotted (with confidence intervals) on the *y* axis and the associated *p*-values on the *x* axis. Values above the red line correspond to a relationship in the direction predicted under the QLE and SSWM approximations. Values to the left of the blue line have *p <* 0.05. Initial condition: *S*(0) = 0.99*N* ^∗^, *I*_00_(0) = 0.01*N* ^∗^, *I*_01_(0) = *I*_10_(0) = *I*_11_(0) = 0.

**Figure S5:**
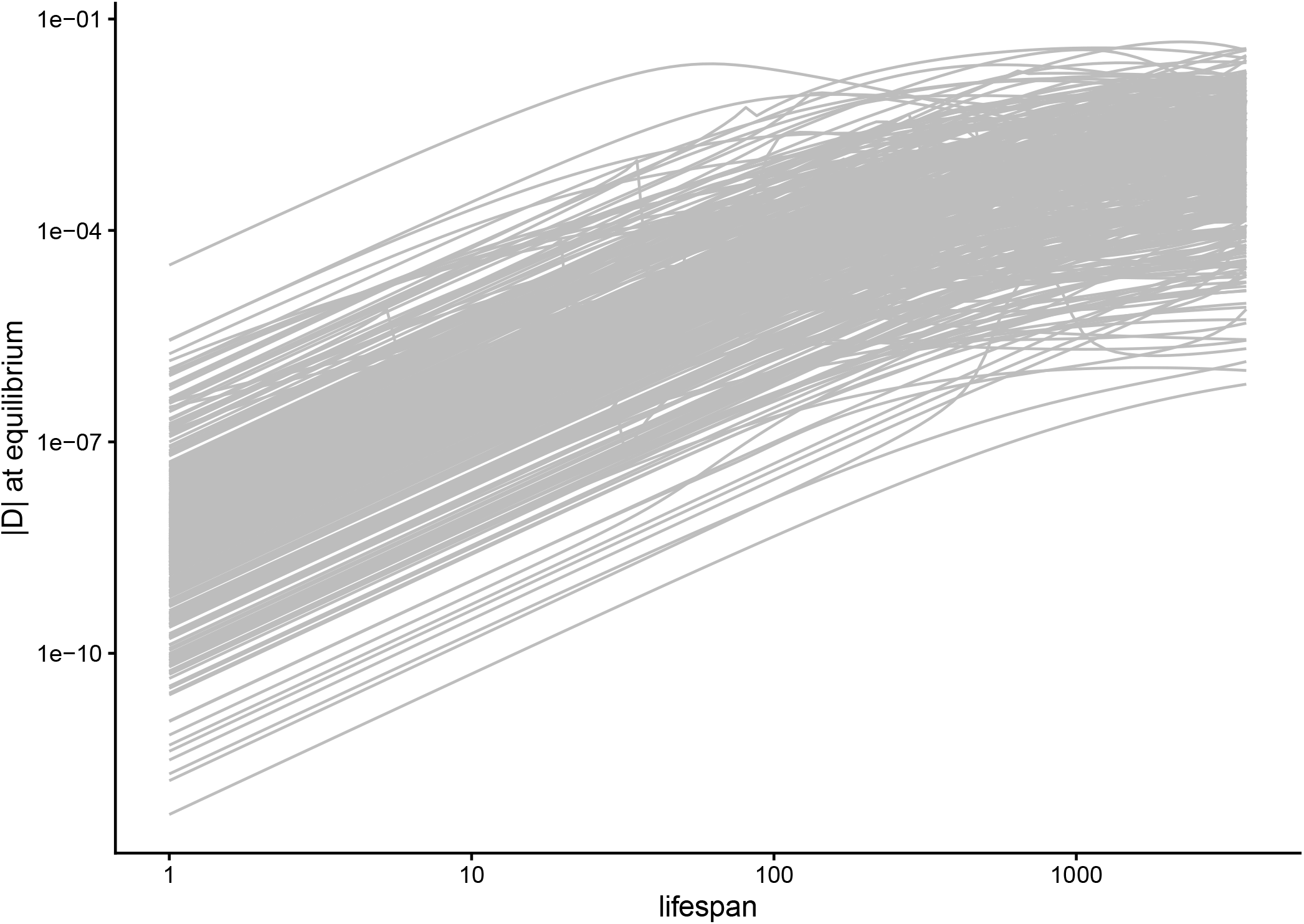
Relationship between |*D*| at equilibrium and host lifespan for all combinations of parameters generated using Latin hypercube sampling. See Figure S4 for more details.

### S4 Analysis of H5N1 genome sequence data

**Figure S6:**
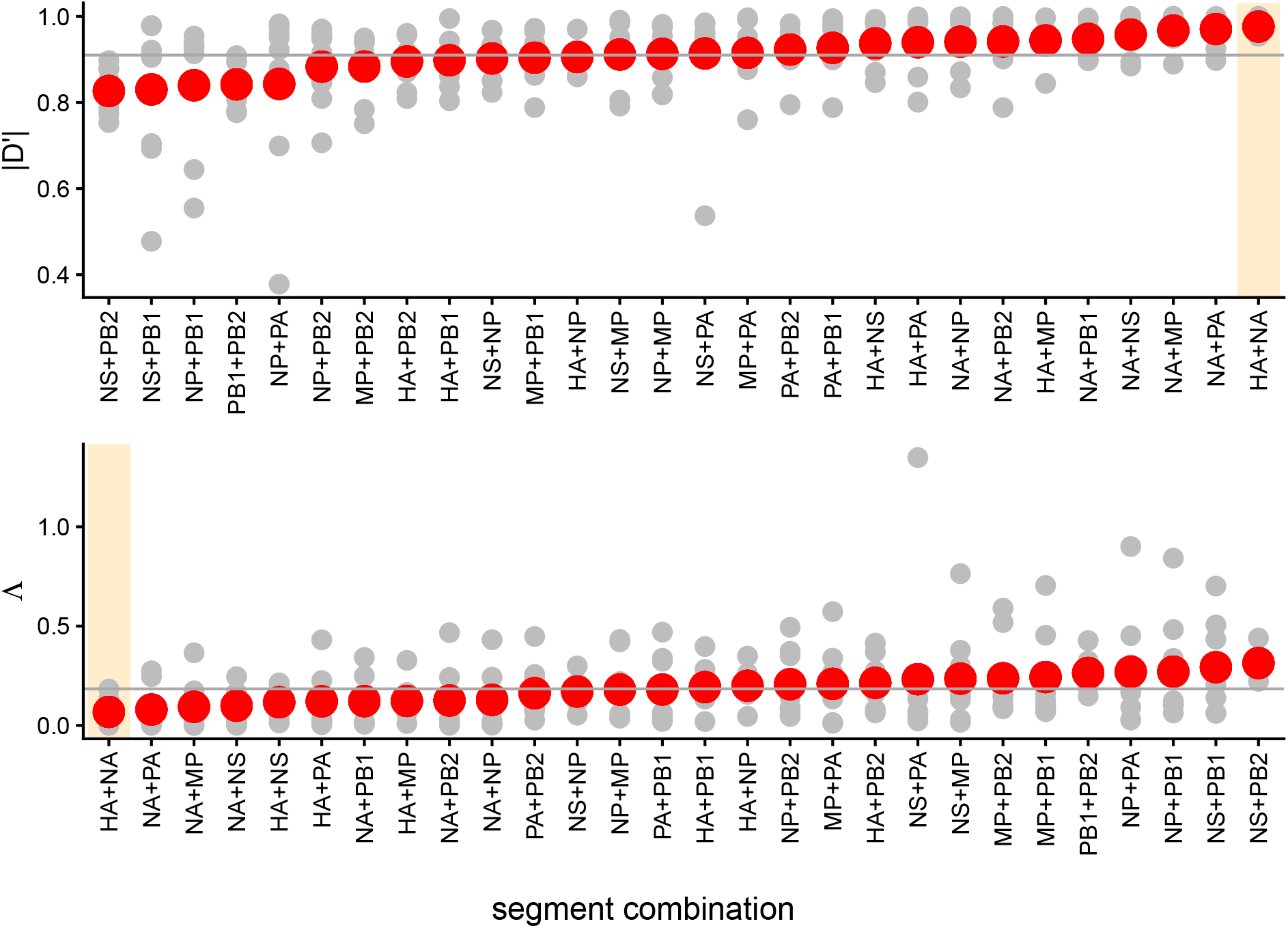
Empirical estimates of |*D*^*′*^| (top row) and Λ (bottom row) from avian influenza data for all segment pairs (HA and NA, PB1 and NP, MP and NA, and so on). Red points are means across all hosts, grey points correspond to the value of a statistic in a given host and segment pair and the grey line denotes the median value over all hosts and segment pairs. The values of *D*^*′*^ (resp., Λ) are highest (resp., lowest) for mutations on the HA and NA (orange band). This is in agreement with what is known about orthogonal functions of these segments and the expectation of epistasis between mutations on them.

**Table S2:** Effect sizes and *p* values for all regressions of *D*^*′*^ and Λ (averaged over mutations on different segments of the flu genome) on host body size, up to 10 significant digits.

| $D'$ | | | | |
| --- | --- | --- | --- | --- |
| | Estimate | Std. Error | $t$ value | $\Pr(>\vert t\vert )$ |
| (Intercept) | 0.8928902530 | 0.0098622675 | 90.5360001108 | 0.0000000000 |
| size | 0.0000097019 | 0.0000046542 | 2.0845506886 | 0.0381252757 |

Table S2: Effect sizes and $p$ values for all regressions of $D'$ and $\Lambda$ (averaged over mutations on different segments of the flu genome) on host body size, up to 10 significant digits.
| $\Lambda$ | | | | |
| --- | --- | --- | --- | --- |
| | Estimate | Std. Error | $t$ value | $\Pr(>\vert t\vert )$ |
| (Intercept) | 0.2187331147 | 0.0195440760 | 11.1917859255 | 0.0000000000 |
| size | -0.0000196615 | 0.0000092232 | -2.1317415255 | 0.0340034696 |

**Table S3:** Effect sizes and *p* values for regression of posterior mean reassortment rates inferred using CoalRe on host body size, up to 10 significant digits.

| | Estimate | Std. Error | $t$ value | $\Pr(>\vert t\vert )$ |
| --- | --- | --- | --- | --- |
| (Intercept) | 0.2291674586 | 0.0172220445 | 13.3066349189 | 0.0000000000 |
| size | -0.0000293949 | 0.0000092986 | -3.1612264638 | 0.0017814137 |

